# Transsynaptic N-cadherin adhesion complexes control presynaptic vesicle and bulk endocytosis at physiological temperature

**DOI:** 10.1101/2021.02.05.429924

**Authors:** Sushma Dagar, Kurt Gottmann

**Affiliations:** Institute of Neuro- and Sensory Physiology, Medical Faculty, Heinrich-Heine-University Düsseldorf, Germany

**Keywords:** N-cadherin, synaptic vesicle endocytosis, bulk endocytosis, physiological temperature, peri-active zone, actin filaments

## Abstract

At mammalian glutamatergic synapses, most basic elements of synaptic transmission have been shown to be modulated by specific transsynaptic adhesion complexes. However, although crucial for synapse homeostasis, a physiological regulation of synaptic vesicle endocytosis by adhesion molecules has not been firmly established. The homophilic adhesion protein N-cadherin is localized at the peri-active zone, where the highly temperature dependent endocytosis of vesicles occurs. Here, we demonstrate an important modulatory role of N-cadherin in endocytosis at near physiological temperature by synaptophysin-pHluorin imaging. Different modes of endocytosis including bulk endocytosis were dependent on N-cadherin expression and function. N-cadherin modulation was mediated by actin filaments, because actin polymerization rescued the knockout induced endocytosis defect. Using super-resolution imaging, we found a strong recruitment of N-cadherin to glutamatergic synapses upon massive vesicle release, which might in turn enhance vesicle endocytosis. This provides a novel, adhesion protein mediated mechanism for efficient coupling of exo- and endocytosis.

## Introduction

A variety of transsynaptic adhesion protein complexes have been molecularly characterized at mammalian CNS synapses, and are thought to have important specific roles in synapse formation, synapse function and synaptic plasticity (Scheiffele et al., 2000; Schreiner et al., 2017; Südhof, 2018). At excitatory glutamatergic synapses, a diversity of synaptic adhesion molecules have been proposed to crucially regulate initial and specific synapse formation (e.g. Sando et al., 2019), active zone and vesicle pool formation (e.g. Han et al., 2018), presynaptic Ca^2+^ channel localization (e.g. Brockhaus et al., 2018), postsynaptic AMPA and NMDA receptor recruitment (e.g. Südhof 2017), and morphological spine plasticity (e.g. Bozdagi et al., 2010). In sharp contrast, no specific molecular adhesion system has been firmly established for transsynaptic regulation of another fundamental aspect of synaptic transmission, physiological synaptic vesicle endocytosis.

The mammalian N-cadherin transsynaptic adhesion complex at glutamatergic synapses is based on pre- and postsynaptic transmembrane N-cadherin proteins containing 5 extracellular cadherin domains (Takeichi, 2007; Benson and Huntley, 2012), which are interacting in *cis* and *trans* configurations (Brasch et al., 2012; Friedman et al., 2015). N-cadherin signaling impacts on the organization of actin filaments in both pre- and postsynaptic compartments via its direct intracellular binding partner beta-catenin and cytoplamic alpha-catenin (Arrikath and Reichardt, 2008; Hirano and Takeichi, 2012). Most importantly, N-cadherin is highly localized at the peri-active zone in the periphery of the synaptic cleft and thus in an optimal strategic position to potentially interfere with actin-dependent vesicle recycling at the endocytic zone (Uchida et al., 1996; Elste and Benson, 2006; Hua et al., 2011; Wu et al., 2016). However, previous studies of N-cadherin functions at the synapse using standard knockout approaches had been difficult, because constitutive N-cadherin knockout mice are lethal at early embryonic stages (Radice et al., 1997), and conditional N-cadherin knockout mice showed strongly impaired neuronal migration during cortex development leading to largely disorganized cortical structures (Kostetskii et al., 2005; Kadowaki et al., 2007).

At the postsynaptic side, a conditional N-cadherin knockout induced after the major period of cortex development revealed an important role of N-cadherin in long-term potentiation (LTP), and a crucial role in LTP-associated morphological spine stabilization in hippocampal neurons *in vivo* (Bozdagi et al., 2010). This study strongly confirmed previous studies using expression of a dominant-negative, function-blocking N-cadherin mutant protein lacking the extracellular cadherin domains (Ncad-ΔE), that had suggested an important function of N-cadherin in postsynaptic spine formation and plasticity (Togashi et al., 2002, Mysore et al., 2008; Mendez et al., 2010; Bian et al., 2015). This purely postsynaptic function of N-cadherin was suggested to be mediated by regulating the spine actin cytoskeleton via catenins (Murase et al., 2002; Abe et al., 2004; Arrikath and Reichardt, 2008; Li et al., 2017). In line with this idea, beta-catenin knockout studies revealed defects in postsynaptic AMPA receptors (Okuda et al., 2007).

In addition to this well established postsynaptic role in spine formation and stabilization, presynaptic functions of the transsynaptic N-cadherin adhesion complex in synaptic vesicle clustering and exocytosis have also been suggested (Bamji et al., 2003; Jüngling et al., 2006, Stan et al., 2010; Vitureira et al., 2011). Subtle defects in short-term plasticity (Jüngling et al., 2006), and in recovery from depletion upon strong stimulation (Jüngling et al., 2006; Vitureira et al., 2011) were demonstrated. In line with the latter findings altered presynaptic vesicle pools have been described upon interfering with beta-catenin expresion and function (Bamji et al., 2003; Chen et al., 2017). Although a few first hints for a potential role of N-cadherin in the regulation of presynaptic vesicle endocytosis have been published (Jüngling et al., 2006; Vitureira et al., 2011; van Stegen et al., 2017), a thorough analysis of synaptic vesicle endocytosis in N-cadherin knockout neurons at close to physiological temperature conditions has not been performed. This is particularly needed, because the modes and kinetics of synaptic vesicle endocytosis are well known to be highly sensitive to temperature (e.g. Granseth and Lagnado, 2008; Soykan et al., 2016; Chanaday and Kavalali, 2018).

In this paper, we have investigated the functional role of N-cadherin in presynaptic vesicle cycling with particular emphasis on vesicle endocytosis at near physiological temperature. We studied vesicle exo- and endocytosis by using synaptophysin-pHluorin fluorescence imaging and by endocytic uptake of fluorescent dyes, thus enabling a sensitive monitoring of synaptic vesicle endocytosis in cultured cortical neurons. In N-cadherin deficient neurons, we observed a strong impairment of vesicle endocytosis at physiological temperature at autaptic and axonal release sites. Both, individual vesicle endocytosis as well as bulk endocytosis were similarly affected by N-cadherin knockout. Mechanistically, signaling of the N-cadherin complex to actin filaments apeared to be crucial as indicated by rescue experiments. Furthermore, we demonstrate a release activity-induced recruitment of N-cadherin to the peri-active zone of glutamatergic synapses by using super resolution microscopy. N-cadherin recruitment might in turn enhance vesicle endocytosis, thus forming the basis of a molecular mechanism for efficient coupling of exo- and endocytosis.

## Results

### SypHy imaging revealed pronounced temperature dependence of synaptic vesicle endocytosis

In this study, we investigated the functional roles of the synaptic adhesion molecule N-cadherin in both synaptic vesicle exo- and endocytosis in cultured mouse cortical neurons. In an initial experimental approach, we used synaptophysin-pHluorin (SypHy) expression to quantitatively study both synaptic vesicle exo- and endocytosis at individual release sites (Granseth et al., 2006). SypHy fluorescence signals at discrete autaptic and axonal puncta (vesicle accumulations; Fig. 1A) increase during synaptic vesicle exocytosis, because of neutralization of the acidic pH of synaptic vesicles, and susequently decrease during synaptic vesicle endocytosis and re-acidification (Royle et al., 2008; Fig. 1B). To reveal all autaptic and axonal puncta expressing SypHy in individual transfected cortical neurons, the pH of synaptic vesicles was neutralized by addition of NH_4_Cl (50 mM) at the end of each experiment (Fig. 1A). To identify SypHy transfected cells, cortical neurons were co-transfectd with DsRed2 visualizing dendrites (MAP2 immuno-positive) and small caliber (tau immuno-positive, MAP2 immuno-negative) axons. SypHy puncta located on dendrites were regarded as autaptic release sites, whereas SypHy puncta located in small caliber axonal processes (without apposed dendrite) were regarded as axonal release sites (Fig. 1A). The SypHy fluorescence signal observed during NH_4_Cl application was used for normalization of the SypHy signals elicited by electrical stimulation at physiological pH at individual autaptic and axonal puncta. Synaptic vesicle exocytosis was quantified by measuring the SypHy fluorescence increase elicited by electrical stimulation at the end of stimulation. Synaptic vesicle endocytosis was quantified by two measures (Fig.1 B): i) by the amount of fluorescence decay at 90 sec after the end of stimulation, and ii) by the monoexponential decay time constant of the SypHy signal.

**Figure 1.**
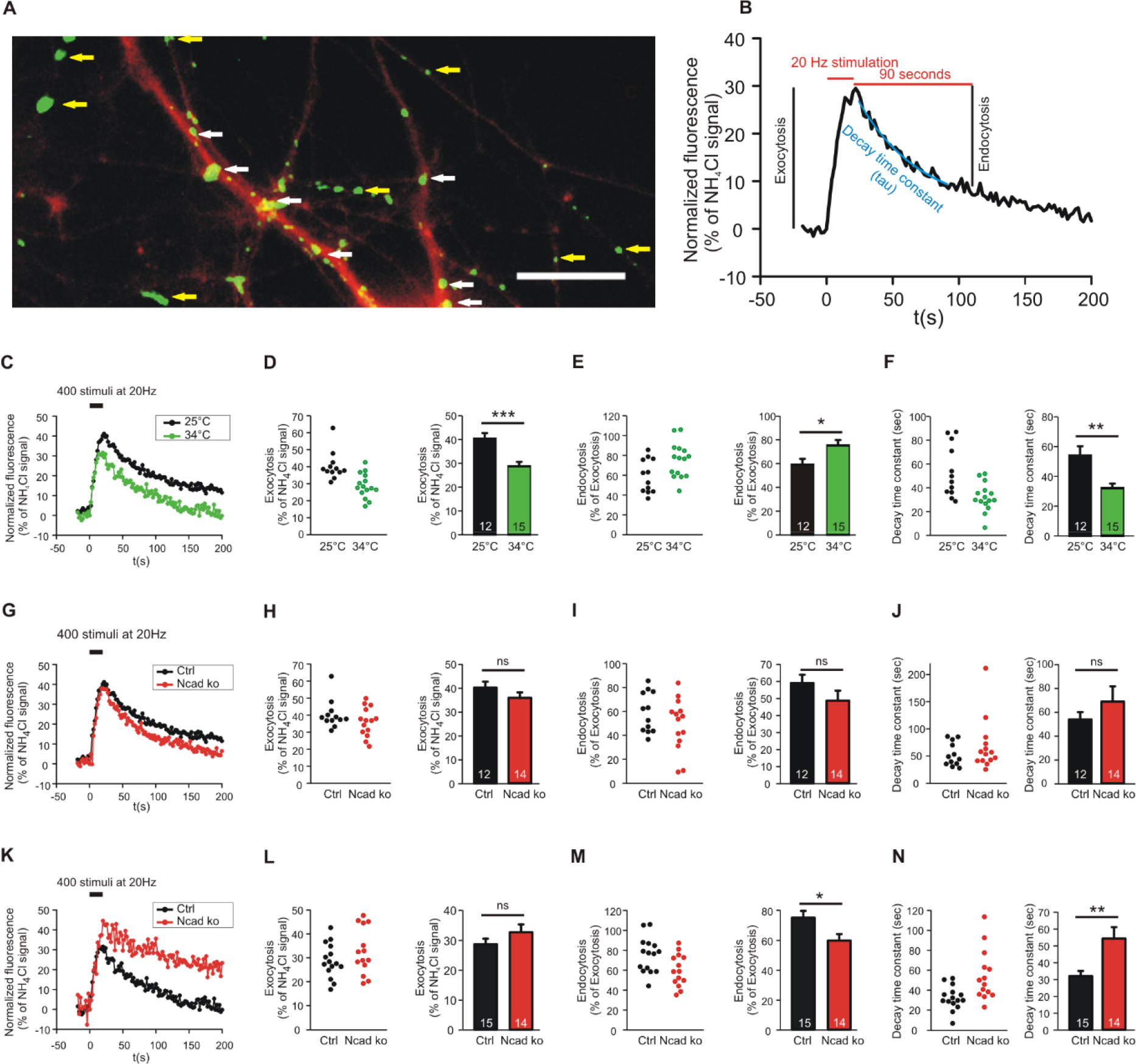
N-cadherin knockout resulted in impaired synaptic vesicle endocytosis at near physiological temperature (34°C) at autapses. **(A)** Example overlay image of cultured cortical neuron processes showing the expression of DsRed2 (transfection marker, red) and the NH_4_Cl-induced SypHy fluorescence (green) at 14-15 DIV. White arrows indicate autapses, and yellow arrows indicate axonal release sites. Scale bar: 10 μm. **(B)** Scheme of quantification of vesicle exocytosis and endocytosis from a SypHy fluorescence signal. Monoexponential fitting (blue) was used to determine decay time constants. **(C-F)** Increasing temperature from room (25°C) to near physiological (34°C) temperature enhanced synaptic vesicle endocytosis at autapses. *n* = 12 (25°C)/15 (34°C) cells, total SypHy puncta analysed were 267/203. **(C)** Example time courses of mean SypHy fluorescence intensities (all puncta per cell averaged, normalized to NH_4_Cl signal) in response to 400 stimuli at 20 Hz at room temperature (25°C, black) and at 34°C (green). **(D)** Quantification of mean SypHy fluorescence intensities at the end of stimulation (20 sec; exocytosis signal) in individual neurons (**left**), and means ± SEM at room and near physiological temperature (**right**). Exocytosis was reduced with enhanced endocytosis kinetics. **(E)** Quantitative analysis of endocytosis at 90 seconds after the end of stimulation (SypHy signal loss as % of exocytosis signal) in individual neurons (**left**), and means ± SEM at room and near physiological temperature (**right**). **(F)** Quantification of SypHy signal decay time constant (tau, monoexponential fit) in individual neurons (**left**), and means ± SEM at room and near physiological temperature (**right**). **(G-J)** Conditional knockout of N-cadherin did not significantly affect synaptic vesicle exo- and endocytosis at room temperature (25°C). *n* = 12 (control)/14 (knockout) cells, total SypHy puncta analysed were 267/268. **(G)** Example time courses of mean SypHy fluorescence intensities (all puncta per cell averaged, normalized to NH_4_Cl signal) for control (co-expressing DsRed2 + SypHy; black) and N-cadherin knockout (co-expressing DsRed2 + SypHy + CreEGFP; red) neurons at 25°C. **(H)** Quantification of mean SypHy fluorescence intensities at the end of stimulation (exocytosis signal) in individual neurons (**left**), and means ± SEM for control (black) and N-cadherin knockout (red) neurons (**right**). **(I)** Quantitative analysis of endocytosis at 90 seconds after the end of stimulation (SypHy signal loss as % of exocytosis signal) in individual neurons (**left**), and means ± SEM for control and N-cadherin knockout neurons (**right**). **(J)** Quantification of SypHy signal decay time constant (tau, monoexponential fit) in individual neurons (**left**), and means ± SEM for control and N-cadherin neurons (**right**). **(K-N)** Conditional knockout of N-cadherin resulted in significantly impaired vesicle endocytosis at near physiological temperature (34°C). *n* = 15 (control)/14 (knockout) cells, total SypHy puncta analysed were 203/268. **(K)** Example time courses of mean SypHy fluorescence intensities (all puncta per cell averaged, normalized to NH_4_Cl signal) for control (black) and N-cadherin knockout (red) neurons at 34°C. **(L)** Quantification of mean SypHy fluorescence intensities at the end of stimulation (exocytosis signal) in individual neurons **(left),** and means ± SEM for control (black) and N-cadherin knockout (red) neurons (**right)**. **(M)** Quantitative analysis of endocytosis at 90 seconds after the end of stimulation (SypHy signal loss as % of exocytosis signal) in individual neurons (**left**), and means ± SEM for control and N-cadherin knockout neurons (**right**). **(N)** Quantification of SypHy signal decay time constant (tau, monoexponential fit) in individual neurons (**left**), and means ± SEMs for control and N-cadherin knockout neurons (**right**). ns: non-significant, * P<0.05, ** P<0.01, *** P<0.001, Student’s *t*-test (E, F, H, I, L, M, N). Whenever normality test failed, Mann-Whitney’s test on ranks was performed (D, J). Data shown in C-F are the same as control data in G-J and K-N. *n* (cells) is indicated on bars.

SypHy imaging was performed in cultured cortical neurons at 14-15 DIV at two different temperatures: i.) at 25°C (room temperature, and ii) at 34°C (near physiological temperature). Both autaptic (Fig. 1C-F) and axonal (Fig. 2A-D) SypHy puncta were analysed separately. As expected, quantitative comparison of the amount of fluorescence decay (endocytosis) at 90 sec revealed a significantly increased amount of endocytosis at 34°C at autaptic release sites (Fig. 1D). Similarly, quantitative comparision of decay time constants revealed a significantly increased rate of endocytosis at 34°C (Fig. 1E). The synaptic vesicle exocytosis related SypHy signal was apparently reduced at 34°C (Fig. 1D), however, this was most likely caused by the enhanced endocytosis ongoing already during the 20 sec stimulation and thus limiting the exocytosis related increase in SypHy fluorescence. At axonal SypHy puncta very similar results were obtained (Fig. 2 A-D). In summary, our results confirm that synaptic vesicle endocytosis is highly temperature sensitive, as has been described also in several previous studies (e.g. Renden and Gersdorff, 2007; Watanabe et al., 2014; Delvendal et al., 2016; Chanaday and Kavalali, 2018).

**Figure 2.**
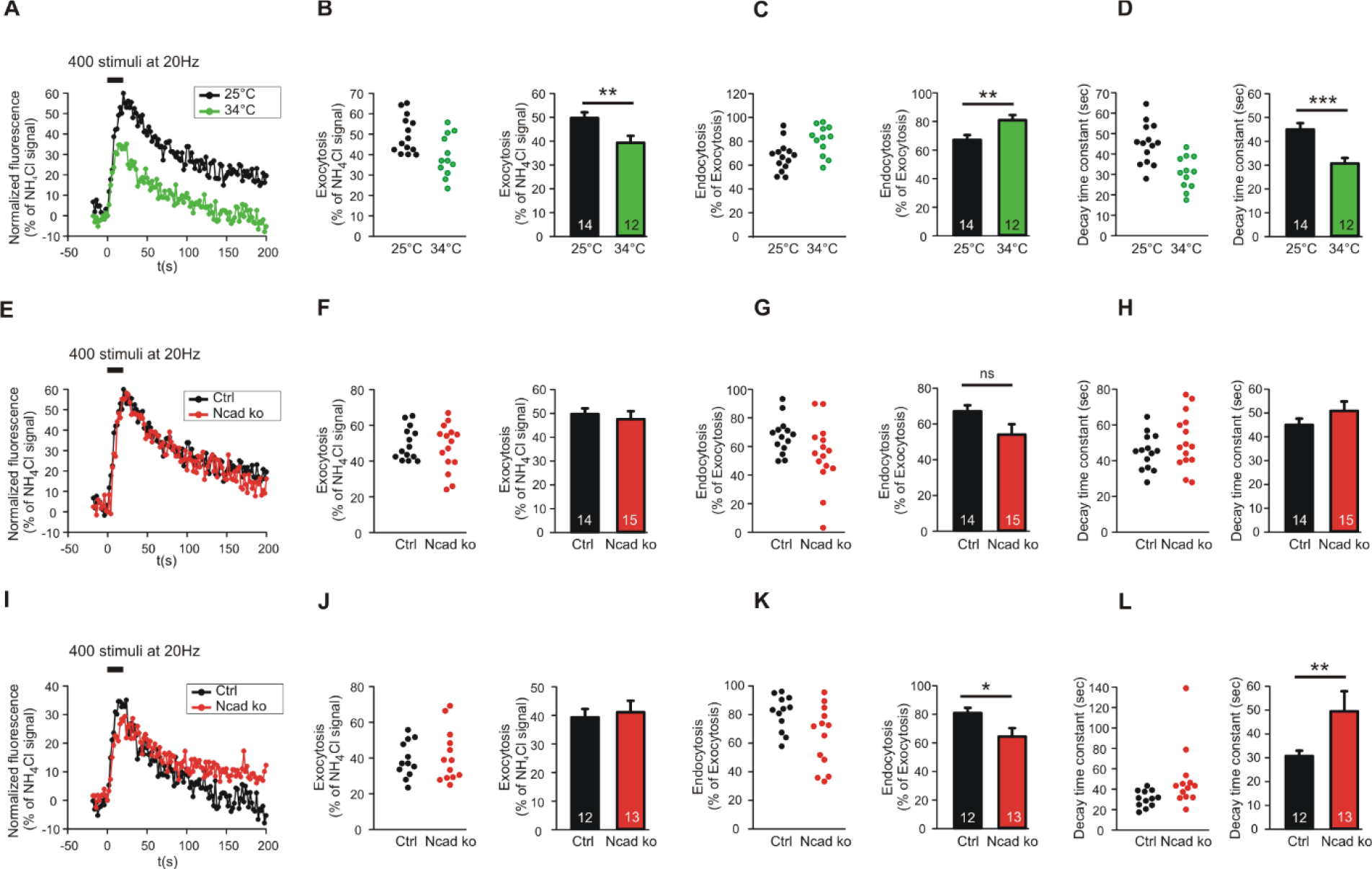
N-cadherin knockout resulted in impaired synaptic vesicle endocytosis at near physiological temperature (34°C) at axonal release sites. **(A-D)** Increasing temperature from room (25°C) to near physiological (34°C) temperature enhanced synaptic vesicle endocytosis at axonal release sites. *n* = 14 (25°C)/12 (34°C) cells, total SypHy puncta analysed were 299/191. **(A)** Example time courses of mean SypHy fluorescence intensities (all puncta per cell averaged, normalized to NH_4_Cl signal) in response to 400 stimuli at 20 Hz at room temperature (25°C, black) and at 34°C (green). **(B)** Quantification of mean SypHy fluorescence intensities at the end of stimulation (20 sec; exocytosis signal) in individual neurons (**left**), and means ± SEM at room and near physiological temperature (**right**). Exocytosis was reduced with enhanced endocytosis kinetics. **(C)** Quantitative analysis of endocytosis at 90 seconds after the end of stimulation (SypHy signal loss as % of exocytosis signal) in individual neurons (**left**), and means ± SEM at room and near physiological temperature (**right**). **(D)** Quantification of SypHy signal decay time constant (tau, monoexponential fit) in individual neurons (**left**), and means ± SEM at room and near physiological temperature (**right**). **(E-H)** Conditional knockout of N-cadherin did not significantly affect synaptic vesicle exo- and endocytosis at room temperature (25°C). *n* = 14 (control)/15 (knockout) cells, total SypHy puncta analysed were 299/370. **(E)** Example time courses of mean SypHy fluorescence intensities (all puncta per cell averaged, normalized to NH_4_Cl signal) for control (co-expressing DsRed2 + SypHy; black) and N-cadherin knockout (co-expressing DsRed2 + SypHy + CreEGFP; red) neurons at 25°C. **(F)** Quantification of mean SypHy fluorescence intensities at the end of stimulation (exocytosis signal) in individual neurons (**left**), and means ± SEM for control (black) and N-cadherin knockout (red) neurons (**right**). **(G)** Quantitative analysis of endocytosis at 90 seconds after the end of stimulation (SypHy signal loss as % of exocytosis signal) in individual neurons (**left**), and means ± SEM for control and N-cadherin knockout neurons (**right**). **(H)** Quantification of SypHy signal decay time constant (tau, monoexponential fit) in individual neurons (**left**), and means ± SEM for control and N-cadherin neurons (**right**). **(I-L)** Conditional knockout of N-cadherin resulted in significantly impaired vesicle endocytosis at near physiological temperature (34°C). *n* = 12 (control)/13 (knockout) cells, total SypHy puncta analysed were 191/237. **(I)** Example time courses of mean SypHy fluorescence intensities (all puncta per cell averaged, normalized to NH_4_Cl signal) for control (black) and N-cadherin knockout (red) neurons at 34°C. **(J)** Quantification of mean SypHy fluorescence intensities at the end of stimulation (exocytosis signal) in individual neurons **(left),** and means ± SEM for control (black) and N-cadherin knockout (red) neurons (**right)**. **(K)** Quantitative analysis of endocytosis at 90 seconds after the end of stimulation (SypHy signal loss as % of exocytosis signal) in individual neurons (**left**), and means ± SEM for control and N-cadherin knockout neurons (**right**). **(L)** Quantification of SypHy signal decay time constant (tau, monoexponential fit) in individual neurons (**left**), and means ± SEMs for control and N-cadherin knockout neurons (**right**). ns: non-significant * P<0.05, ** P<0.01, *** P<0.001, Student’s *t*-test (B, C, D, F, G, H, J, K). Whenever normality test failed, Mann-Whitney’s test on ranks was performed (L). Data shown in A-D are the same as control data in E-H and I-L. *n* (cells) is indicated on bars.

### N-cadherin knockout resulted in impaired endocytosis at near physiological temperature

To study the potential role of N-cadherin in synaptic vesicle exo- and endocytosis, we induced a conditional N-cadherin knockout in individual cortical neurons (with floxed N-cadherin gene; Kostetskii et al., 2005) in culture by sparse co-transfection with cre and SypHy (+ DsRed2). At room temperature (25°C) with application of 400 stimuli at 20 Hz, SypHy imaging at autaptic release sites revealed only a non-significant trend to a reduction in the amount of endocytosis at 90 sec after the end of stimulation in N-cadherin knockout neurons (pre- and postsynaptic knockout) (Fig. 1G, I). Similarly, the SypHy signal decay time constant showed only a non-significant trend to be increased (Fig. 1J). The exocytosis related SypHy signal was not significantly altered in N-cadherin knockout neurons (Fig. 1H). At axonal SypHy puncta very similar results were obtained (Fig. 2E-H), despite the fact that only the presynaptic cell was a N-cadherin knockout neuron. Most importantly, at 34°C with a near physiological rate of endocytosis SypHy imaging at autaptic release sites clearly demonstrated a significant reduction in the amount of endocytosis at 90 sec after the end of stimulation in N-cadherin knockout neurons (pre- and postsynaptic knockout) (Fig. 1K, M). In addition, the SypHy signal decay time constant was significantly increased (Fig. 1N). Again, the exocytosis related SypHy signal was not significantly altered in N-cadherin knockout neurons (Fig. 1L). At axonal SypHy puncta (only presynaptic knockout) a similar significant impairment of endocytosis was observed (Fig. 2I-L). In summary, our results with conditional N-cadherin knockout experiments indicate that N-cadherin plays an important role in enabling highly efficient endocytosis at near physiological temperature. At room temperature endocytosis occurs in a much less efficient way and consequently the effects of N-cadherin deficiency were much less pronounced.

### N-cadherin deficiency impaired both single vesicle endocytosis and bulk endocytosis

Synaptic vesicle endocytosis is well known to occur in at least two, at the subcellular level clearly distinct modes: i) endocytosis of individual synaptic vesicles, and ii) bulk endocytosis of large membrane invaginations. We next wanted to investigate whether individual vesicle endocytosis and bulk endocytosis are both modulated by N-cadherin. We decided to study this by using two rather different experimental conditions: i) a stimulation paradigm under which bulk endocytosis is almost completely absent, and ii) a second stimulation paradigm under which bulk endocytosis is prominent. To specifically label bulk endocytosis events, we stimulated the neurons in the presence of extracellular fluorescent TMR-dextran (40kD), because it is well established to be taken up selectively into large membrane invaginations (Clayton and Cousin, 2009b). To unspecifically visualize all endocytosis events, we co-added the styryl dye FM1-43 (Gaffield and Betz, 2006) to the extacellular solution. We reasoned that individual vesicle endocytosis events will be labeled only by FM1-43, whereas bulk endocytosis events will be labeled by both FM1-43 and TMR-dextran (Clayton and Cousin, 2009b). We initially used two different electrical stimulation paradigms (200 stimuli and 400 stimuli at 20Hz), which were each applied at two different temperatures (25°C and 34°C) to mass cultures of cortical neurons at 12-13 DIV (Fig. 3 A, B). We observed that with 200 stimuli at 25°C only rather few FM1-43 puncta were co-labeled with TMR-dextran (Fig. 3C) thus representing a stimulation condition where bulk endocytosis is largely absent. In contrast, with 400 stimuli at 34°C more than 50% of the FM1-43 puncta were co-labeled with TMR-dextran (Fig. 3C) thus representing a stimulation condition where bulk endocytosis is prominent.

**Figure 3.**
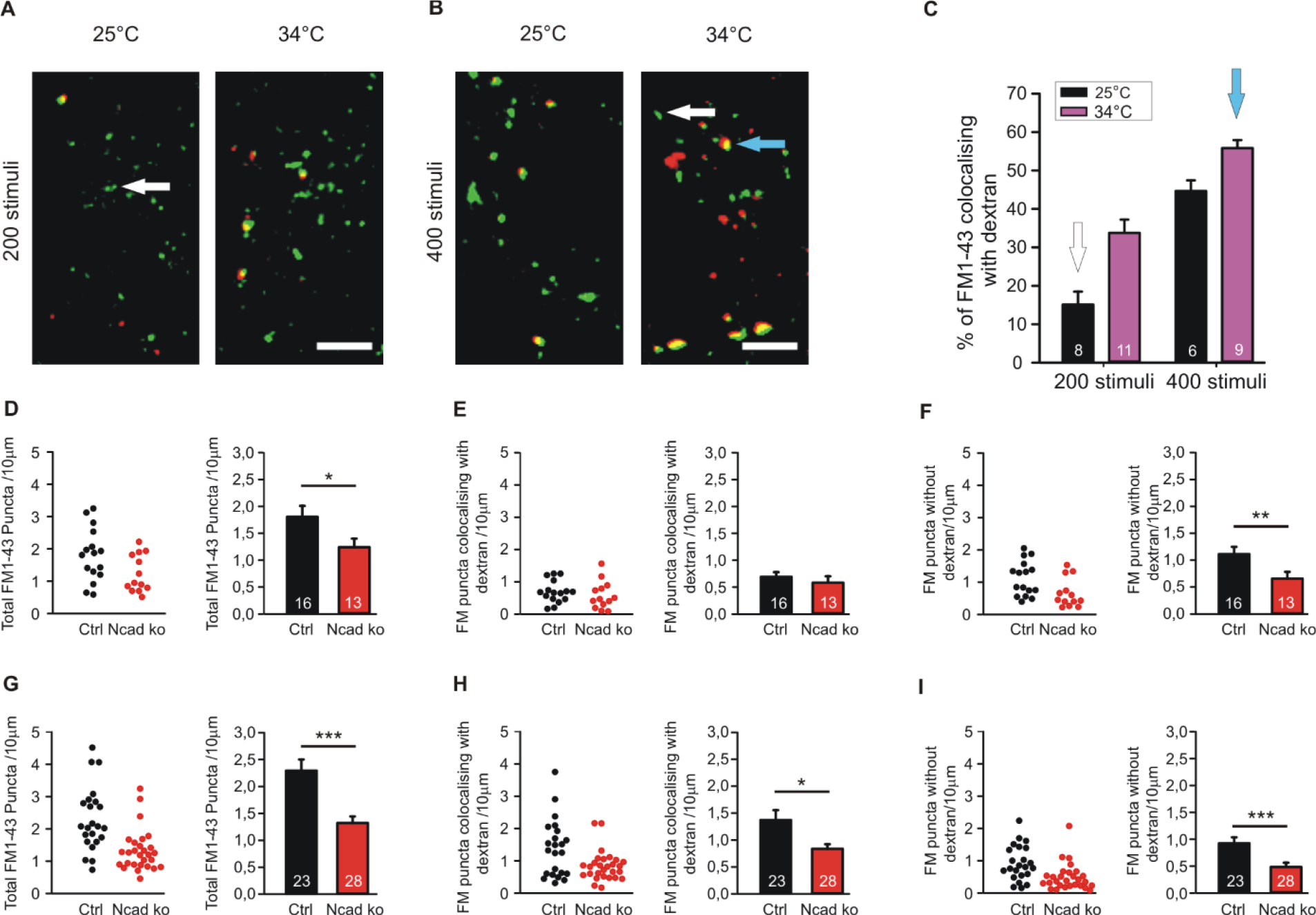
Knockout of N-cadherin resulted in impairment of different modes of endocytosis at near physiological temperature (34°C). (**A-C**) Experimental paradigm to distinguish between individual vesicle endocytosis and bulk endocytosis. (**A**) Representative overlay images showing FM1-43 (green puncta) and TMR-dextran (red puncta) uptake induced by 200 stimuli (at 20 Hz) in cortical neurons in mass culture at 25°C (**left**) and at 34°C (**right**). White arrow indicates a typical synaptic punctum with only FM1-43 uptake. Scale bar: 5 μm. **(B)** Example overlay images showing FM1-43 (green puncta) and TMR-dextran (red puncta) uptake induced by 400 stimuli (at 20 Hz) in cortical neurons in mass culture at 25°C (**left**) and at 34°C (**right**). Blue arrow indicates a typical punctum (yellow) showing co-uptake of both FM1-43 and TMR-dextran. Scale bar: 5 μm. **(C)** Quantification of the percentage of FM1-43 positive puncta co-localizing with TMR-dextran uptake (black 25°C; magenta 34°C) with 200 (*n* = 8/11) and 400 stimuli (*n* = 6/9, *n* represents fields of view). **(D-F)** Cre mediated conditional (postsynaptic) knockout of N-cadherin resulted in reduced individual vesicle endocytosis at 25°C in cortical neurons at 14-15 DIV. Cortical neurons were sparsely co-transfected at 9 DIV with EBFP2 (to visualize dendrites) and CreEGFP. FM1-43 and TMR-dextran uptake was induced by electrical stimulation (200 stimuli at 20 Hz at 25°C). **(D)** Quantification of the dendritic density of total FM1-43 labeled puncta in individual transfected neurons (**left**) and means ± SEM (black: control; red: N-cadherin knockout; *n* = 16/13 cells, **right**). **(E)** Quantification of the dendritic density of FM1-43 puncta co-localizing with TMR-dextran in individual neurons (**left**) and means ± SEM (**right**). Note that dextran uptake was very weak at this experimental condition (baseline uptake). **(F)** Quantification of the dendritic density of FM1-43 puncta not co-localizing with TMR-dextran in individual neurons (**left**) and means ± SEM (**right**). **(G-I)** Postsynaptic knockout of N-cadherin by sparse Cre transfection (+EBFP2) resulted in a significant reduction in both, individual vesicle endocytosis and bulk endocytosis at near physiological temperature (34°C). Vesicle cycling was induced by 400 stimuli at 20 Hz. **(G)** Quantification of the dendritic density of total FM1-43 puncta in individual neurons (**left**) and means ± SEM (black: control; red: N-cadherin knockout; *n* = 23/28 cells, **right**). **(H)** Quantification of the dendritic density of FM1-43 puncta co-localizing with TMR-dextran in individual neurons **(left)** and means ± SEM (**right**). Note the reduction in TMR-dextran uptake in N-cadherin knockout neurons indicating impaired bulk endocytosis. **(I)** Quantification of the dendritic density of FM1-43 puncta not co-localizing with TMR-dextran in individual neurons (**left**) and means ± SEM (**right**). * P<0.05, ** P<0.01, *** P<0.001, Student’s *t*-test (D, E). Whenever normality test failed Mann-Whitney’s test on ranks was performed (F, G, H, I).

After having established these two distinctive stimulation paradigms, we studied FM1-43 and TMR-dextran uptake in individual cortical neurons (with floxed N-cadherin genes) that were visualized by EBFP2 expression (transfection at 9 DIV) at 14-15 DIV in mass culture. Co-expression of cre was used to induce a conditional N-cadherin knockout in individual transfected neurons (postsynaptic knockout). With bulk endocytosis (and TMR-dextran uptake) largely absent (200 stimuli at 25°C), we observed a significant decrease in the dendritic density of FM1-43 puncta in N-cadherin deficient neurons (Fig. 3D, F). This indicates an important dependence of individual vesicle endocytosis on N-cadherin expression. With bulk endocytosis occurring in addition (400 stimuli at 34°C), we again observed a significant reduction in the dendritic density of FM1-43 puncta in N-cadherin knockout neurons (Fig. 3G). The selective analysis of FM1-43 puncta that were co-labeled with TMR-dextran revealed a significant decrease of the dendritic density of FM1-43 puncta co-localizing with TMR-dextran (Fig. 3H). This indicates a strong dependence also of bulk endocytosis on N-cadherin expression. The selective analysis of FM1-43 puncta that were not co-labeled again revealed a significant reduction of the dendritic density of FM1-43 only puncta (individual vesicle endocytosis, Fig. 3I). In summary, our experiments analysing the N-cadherin dependence of FM1-43 and TMR-dextran uptake revealed that both individual vesicle endocytosis and bulk endocytosis are impaired in N-cadherin deficient neurons.

### Inhibition of N-cadherin function resulted in impaired endocytosis at near physiological temperature

We next wanted to study, whether an inhibition of N-cadherin function leaving the N-cadherin gene intact has similar effects on synaptic vesicle endocytosis as the complete knockout of the N-cadherin gene. Overexpression of a transmembrane C-terminal fragment of N-cadherin without the extracellular cadherin domains (NcadΔE) is a well established experimental approach to inhibit N-cadherin function (Togashi et al., 2002; Andreyeva et al., 2012). We first investigated the effects of NcadΔE overexpression on synaptic vesicle endocytosis by using SypHy imaging in microisland cultures of mouse cortical neurons at near physiological temperature (34°C). At 9 DIV individual neurons were co-transfected with a transfection marker (DsRed2), a SypHy vector, and a NcadΔE vector. At 12-13 DIV SypHy fluorescence signals were elicited by extracellular stimulation (400 stimuli at 20 Hz), and SypHy signals from autaptic puncta were analysed as described above (Fig. 4A). In NcadΔE overexpressing neurons we found a significant reduction in the fraction of endocytosis at 90 sec after the end of stimulation (Fig. 4C), and a significant increase in the decay time constant of the SypHy signal (Fig. 4D). The exocytosis related SypHy signal was not altered (Fig. 4B). These results demonstrate that inhibition of N-cadherin function results in an impaired synaptic vesicle endocytosis.

**Figure 4.**
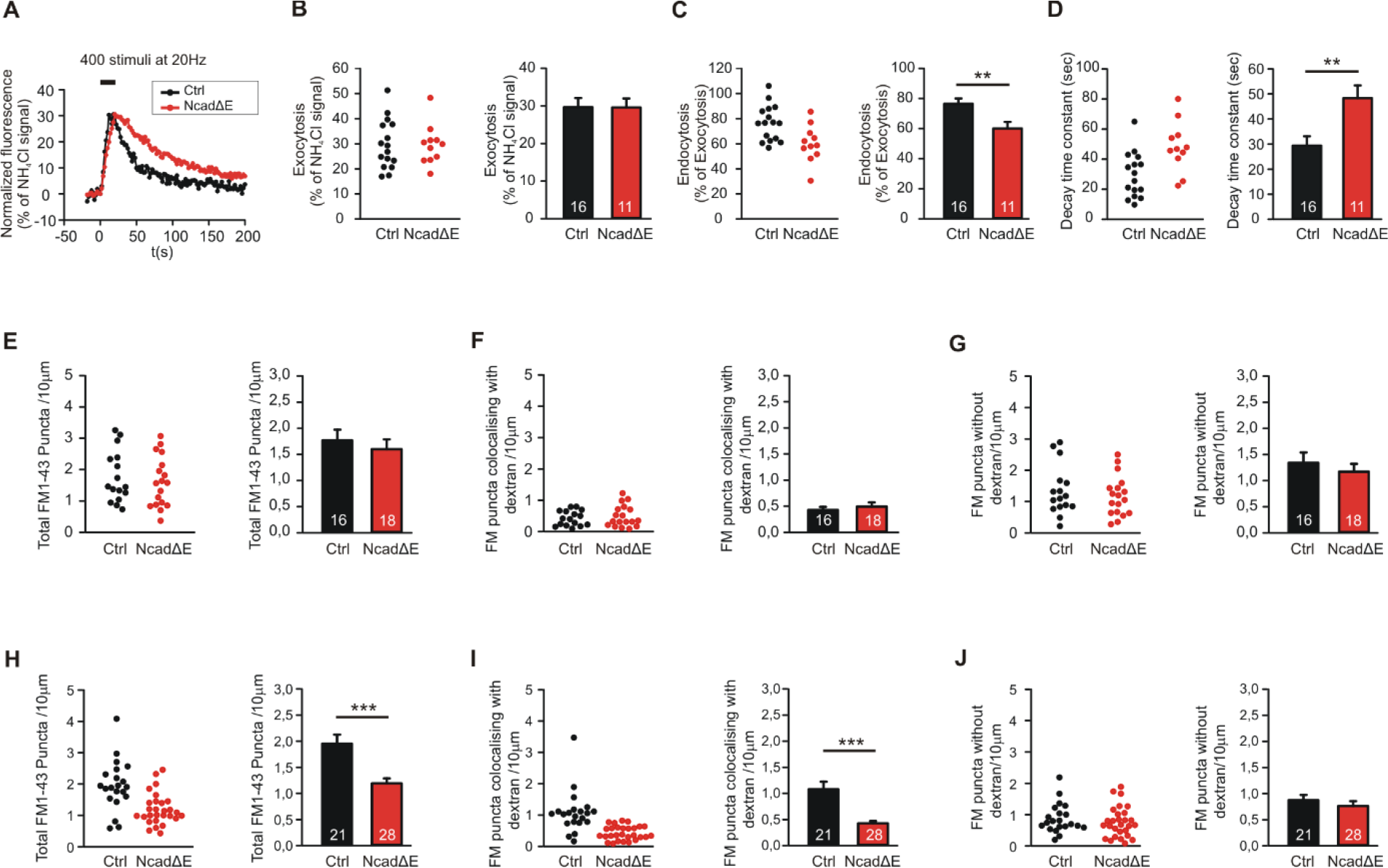
Inhibition of N-cadherin function by NcadΔE expression resulted in reduced synaptic vesicle endocytosis. **(A-D)** Inhibiting the function of N-cadherin by co-expressing a dominant-negative N-cadherin fragment (NcadΔE) lacking the extracellular domains (with DsRed2 and SypHy at 9 DIV in individual cortical neurons) resulted in a significant reduction in synaptic vesicle endocytosis at autapses at near physiological temperature (34°C). SypHy imaging was performed at 12-14 DIV with 400 stimuli at 20 Hz. *n*=16 (control)/11 (NcadΔE) cells, total number of SypHy puncta analysed were 463/192. **(A)** Example time courses of mean SypHy fluorescence intensities (all puncta per cell averaged, normalized to NH_4_Cl signal) for control (co-expressing DsRed2 + SypHy; black) and NcadΔE expressing (co-expressing DsRed2 + SypHy + NcadΔE; red) neurons at 34°C. **(B)** Quantification of mean SypHy fluorescence intensities at the end of stimulation (20 s; exocytosis signal) in individual neurons (**left**), and means ± SEM (**right**) for control (black) and NcadΔE expressing (red) neurons. **(C)** Quantitative analysis of endocytosis at 90 seconds after the end of stimulation (SypHy signal loss as % of exocytosis signal) in individual neurons (**left**), and means ± SEM for control and N-cadΔE expressing neurons (**right**). **(D)** Quantification of SypHy signal decay time constant (tau, monoexponential fit) in individual neurons (**left**), and means ± SEM for control and N-cadΔE expressing neurons (**right**). **(E-G)** Postsynaptic functional block of N-cadherin by sparse NcadΔE expression in mass cultures of cortical neurons did not result in a significant change in vesicle endocytosis at 25°C, as studied by uptake of FM1-43 and TMR-dextran induced by 200 stimuli at 20 Hz. **(E)** Quantification of the dendritic density of total FM1-43 puncta in individual transfected neurons (**left**), and means ± SEM (black: control; red: N-cadΔE expression; *n*=16/18 cells, **right**). **(F)** Quantification of the dendritic density of FM1-43 puncta co-localizing with TMR-dextran in individual neurons (**left**), and means ± SEM (**right**). TMR-dextran uptake is very low at this experimental condition (baseline uptake). **(G)** Quantification of the dendritic density of FM1-43 puncta not co-localizing with TMR-dextran in individual neurons (**left**), and means ± SEM (**right**). **(H-J)** Postsynaptic inhibition of N-cadherin function by NcadΔE expression resulted in a significant impairment in bulk endocytosis at 34°C (400 stimuli at 20 Hz). **(H)** Quantification of the dendritic density of total FM1-43 puncta in individual transfected neurons (**left**), and means ± SEM (black: control; red: NcadΔE expression; *n*=21/28 cells, **right**). **(I)** Quantification of the dendritic density of FM1-43 puncta co-localizing with TMR-dextran in individual neurons (**left**), and means ± SEM (**right**). **(J)** Quantification of the dendritic density of FM1-43 puncta not co-localizing with TMR-dextran in individual neurons (**left**), and means ± SEM (**right**). ** P<0.01, *** P<0.001, Student’s *t*-test (B-D). Whenever normality test failed Mann-Whitney’s test on ranks was performed (E-J).

We further investigated the effects of NcadΔE overexpression on the endocytotic uptake of FM1-43 and TMR-dextran, which was studied as described above. In line with inhibition of N-cadherin function being a somewhat weaker interference than N-cadherin gene knockout, we did not observe a significant reduction of FM1-43 uptake (vesicle endocytosis) at 25°C upon NcadΔE overexpression (Fig. 4E-G). In contrast, at a near physiological temperature (34°C) we found a significant decrease in the dendritic density of FM1-43 puncta (Fig. 4H). This was mainly due to a significant reduction of FM1-43 puncta co-localizing with TMR-dextran (Fig. 4I), whereas the dendritic density of FM1-43 puncta not co-localizing with TMR-dextran did not change (Fig. 4J). These results indicate that partial inhibition of N-cadherin function results in an impairment of bulk endocytosis, whereas direct individual vesicle endocytosis is largely maintained. This is in line with the idea that bulk endocytosis is a more complex process than direct vesicle endocytosis, and therefore is more dependent on full N-cadherin function, which potentially controls the complexity of actin organization.

### N-cadherin knockout additionally resulted in an impaired exocytosis of synaptic vesicles

Based on patch clamp recordings, it had been described previously by our group, that a N-cadherin knockout in mouse ES cell-derived neurons resulted in a defect in exocytosis of synaptic vesicles when a two pulse stimulation paradigm was used (Jüngling et al., 2006). However, in the above described SypHy imaging experiments we did not observe a significantly reduced exocytosis signal when using a relatively long lasting stimulation paradigm (400 stimuli at 20 Hz). Differences in exocytosis might have been obscured with this stimulation paradigm by the fact that endocytosis already limits SypHy signal amplitude during the 20 sec stimulation. Therefore, we further studied synaptic vesicle exocytosis with SypHy imaging with a shortened stimulation paradigm (100 stimuli at 20 Hz). Using this 5 sec stimulation and identical transfection and cultivation conditions as described above, we indeed found a significant reduction of the exocytosis-related SypHy signal amplitude at autaptic sites in N-cadherin knockout neurons (Fig. 5A, B). The analysis of axonal sites revealed a similar reduction of the exocytosis-related SypHy signal in N-cadherin deficient neurons (Fig. 5C, D). Despite the weaker stimulation, the analysis of the endocytosis-related SypHy signal decay revealed a similar tendency towards an impairment of endocytosis in N-cadherin knockout neurons (Suppl.Fig. 1) as found above with the 400 stimuli at 20 Hz stimulation paradigm.

**Figure 5.**
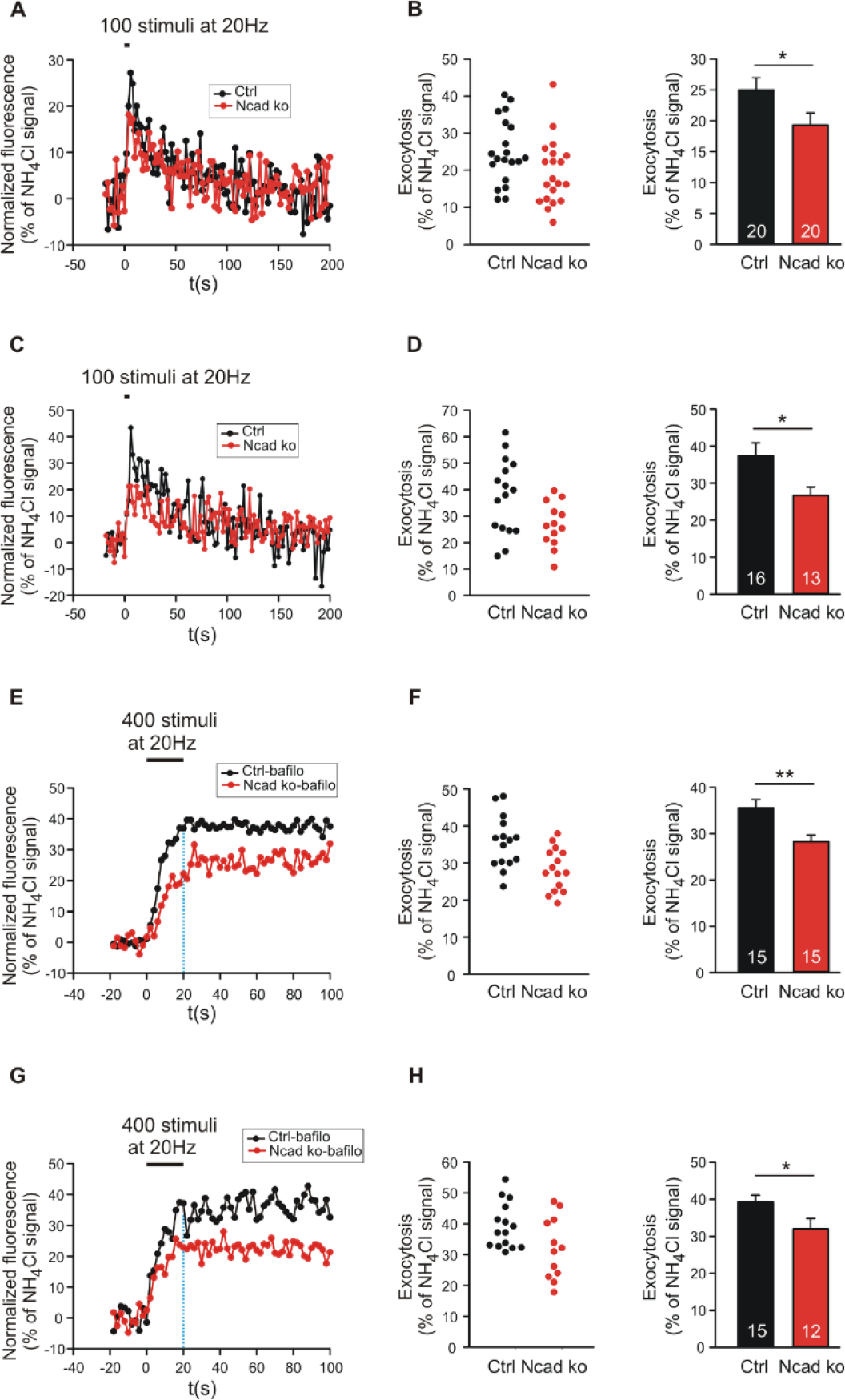
Knockout of N-cadherin resulted in reduced synaptic vesicle exocytosis with short term stimulation at near physiological temperature (34°C). (**A, B**) Effect of N-cadherin knockout on synaptic vesicle exocytosis studied with short term stimulation at autapses in cultured cortical neurons. (**A**) Example time courses of mean SypHy fluorescence intensities (all puncta per cell averaged, normalized to NH_4_Cl signal) for control (co-expressing DsRed2 + SypHy; black) and N-cadherin knockout (co-expressing DsRed2 + SypHy + CreEGFP; red) neurons in response to 100 stimuli at 20 Hz at 34°C. (**B**) Quantification of mean SypHy intensities at the end of stimulation (5 s; exocytosis signal) in individual neurons (**left**), and means ± SEM (**right**) for control (black) and N-cadherin knockout (red) neurons. *n* = 20/20 cells, total SypHy puncta analysed were 333/383. **(C, D)** Effect of N-cadherin knockout on synaptic vesicle exocytosis studied with short term stimulation at axonal release sites. (**C**) Example time courses of mean SypHy fluorescence intensities (all puncta per cell averaged, normalized to NH_4_Cl signal) for control (black) and N-cadherin knockout (red) neurons in response to 100 stimuli at 20 Hz at 34°C. (**D**) Quantification of mean SypHy intensities at the end of stimulation (5 s; exocytosis signal) in individual neurons (**left**), and means ± SEM (**right**) for control (black) and N-cadherin knockout (red) neurons. *n* = 16/13 cells, total SypHy puncta analysed were 212/130. Student’s *t*-test was used for statistical analysis,* P<0.05 (B, D). (**E, F**) Effect of N-cadherin knockout on synaptic vesicle exocytosis studied in the presence of bafilomycin-A1 at autapses. (**E**) Example time courses of mean SypHy fluorescence intensities (all puncta per cell averaged, normalized to NH_4_Cl signal) for control (black) and N-cadherin knockout (red) neurons in response to 400 stimuli at 20 Hz at 34°C in the presence of bafilomycin-A1 (1 μM). (**F**) Quantification of mean SypHy intensities at the end of stimulation (20 s; exocytosis signal) in individual neurons (**left**), and means ± SEM (**right**) for control (black) and N-cadherin knockout (red) neurons. *n* = 15/15 cells, total SypHy puncta analysed were 361/403. (**G, H**) Effect of N-cadherin knockout on synaptic vesicle exocytosis studied in the presence of bafilomycin-A1 at axonal release sites. (**G**) Example time courses of mean SypHy fluorescence intensities (all puncta per cell averaged, normalized to NH_4_Cl signal) for control (black) and N-cadherin knockout (red) neurons in response to 400 stimuli at 20 Hz at 34°C in the presence of bafilomycin-A1 (1 μM). Dotted blue line indicates the end of stimulation. (**H**) Quantification of mean SypHy intensities at the end of stimulation (20 s; exocytosis signal) in individual neurons (**left**), and means ± SEM (**right**) for control (black) and N-cadherin knockout (red) neurons. *n* = 15/12 cells, total SypHy puncta analysed were 289/275. Student’s *t*-test was used for statistical analysis, * P<0.05, ** P<0.01.

To further confirm changes in synaptic vesicle exocytosis in N-cadherin deficient cortical neurons, we performed SypHy imaging in the presence of bafilomycin A1 (1 μM). Bafilomycin A1 blocks the reacidifiction of newly endocytosed vesicles/membrane invaginations (Atluri and Ryan, 2006) thus leading to a non-decaying fluorescence increase upon stimulation that represents the exocytosis of synaptic vesicles (Fig. 5E, G). As expected, we observed a significant reduction of the SypHy fluorescence signal at the end of a 20 sec stimulation (400 stimuli at 20 Hz) at both autaptic and axonal SypHy puncta in N-cadherin knockout neurons (Fig. 5 F, H). In summary, our results indicate that N-cadherin plays a significant role also in synaptic vesicle exocytosis, possibly by contributing via modulating the actin cytoskeleton to release site clearance after vesicle fusion.

### Enhancing actin polymerization rescued endocytosis defects in N-cadherin knockout neurons

To address the molecular mechanisms underlying the enhancing role of N-cadherin in synaptic vesicle endocytosis, we studied whether promoting actin polymerization can rescue the endocytosis defects observed in N-cadherin knockout neurons. We used the pharmacological drug jasplakinolide to promote actin polymerization in cultured neurons (Soykan et al., 2017), and performed SypHy imaging at near physiological temperature (34°C) with standard stimulation (400 stimuli at 20 Hz). Acute (5 min) extracellular addition of jasplakinolide (5 μM) did not significantly alter the exoytosis-related as well as the endocytosis-related SypHy signal components at autaptic release sites in control cortical neurons (Fig. 6). Importantly, in N-cadherin knockout neurons acute addition of jaspaklinolide completely rescued the slow down of endocytosis kinetics that was induced by the N-cadherin deficiency (Fig. 6A, D). Furthermore, a clear trend to an increased amount of endocytosis at 90 sec after the end of stimulation was observed in N-cadherin knockout neurons upon acute addition of jasplakinolide (Fig. 6A, C). These results suggest that N-cadherin modulates endocytosis by its well known signaling to the actin network via β- and α-catenins thus leading to an improved organization of actin filaments.

**Figure 6.**
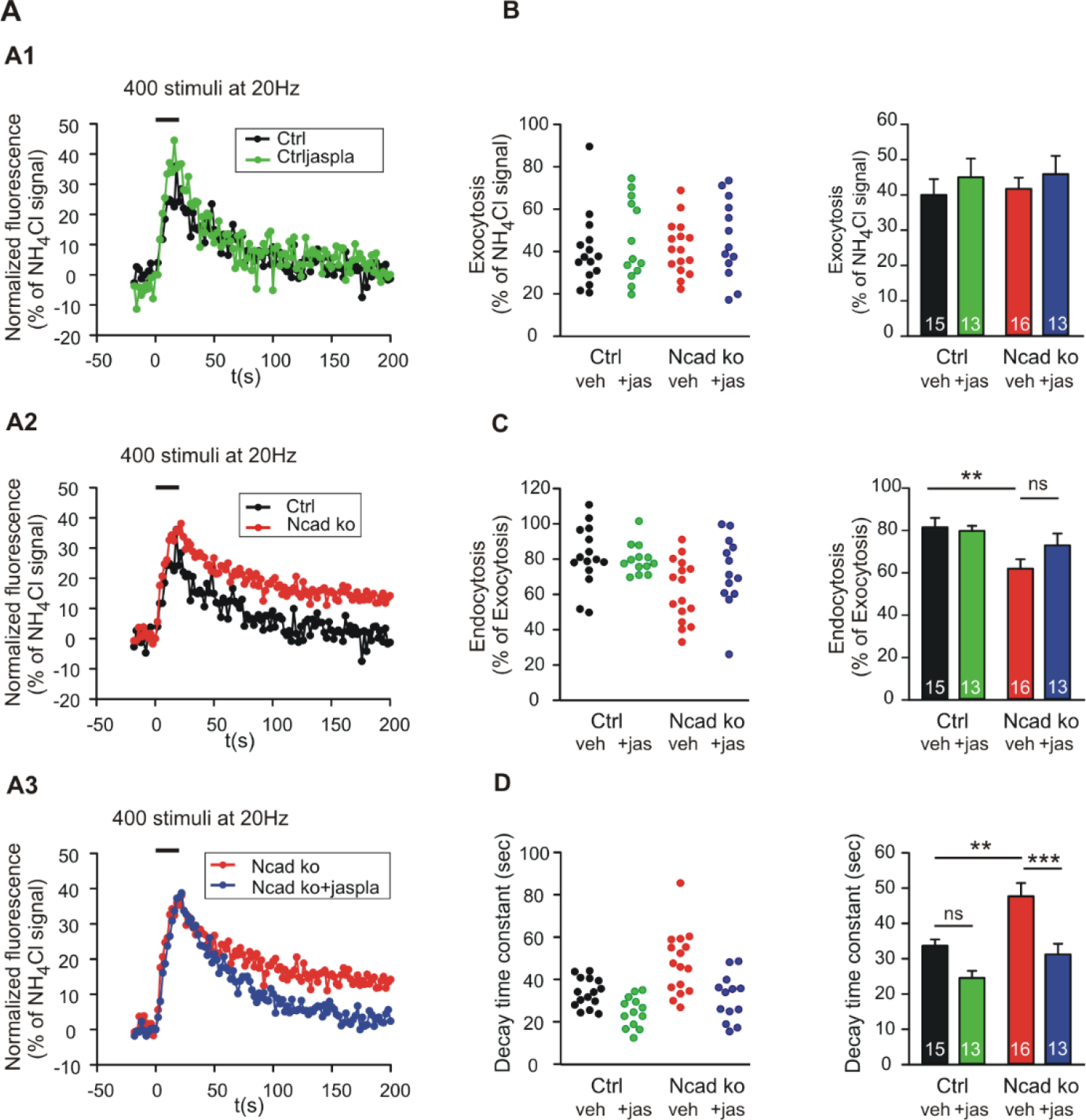
Polymerization of actin induced by Jasplakinolide in N-cadherin knockout neurons rescued impaired synaptic vesicle endocytosis. **(A-D)** SypHy imaging was performed at autapses (400 stimuli at 20 Hz at 34°C) in control and N-cadherin knockout neurons at 14-15 DIV (see Fig. 1). Cultures were treated with 5 μM Jasplakinolide for 5 min before imaging to induce actin polymerization. **(A1)** Example time courses of mean SypHy fluorescence intensities (all puncta per cell averaged, normalized to NH_4_Cl signal) for control neurons (co-expressing DsRed2 + SypHy; black) and control neurons treated with 5 μM Jasplakinolide (green). **(A2)** Example time courses of mean SypHy fluorescence intensities for control neurons (black) and N-cadherin knockout neurons (co-expressing DsRed2 + SypHy + CreEGFP, red). **(A3)** Example time courses of mean SypHy fluorescence intensities for N-cadherin knockout neurons (red) and N-cadherin knockout neurons treated with 5 μM Jasplakinolide (blue). Note that SypHy traces for control (A1, A2) and N-cadherin knockout (A2, A3) neurons are shown twice for clarity. **(B)** Quantification of mean SypHy intensities at the end of stimulation (20 s; exocytosis signal) in individual neurons (**left**), and means ± SEM (**right**) for control neurons (black), control neurons treated with Jasplakinolide (green), N-cadherin knockout neurons (red), and N-cadherin knockout neurons treated with Jasplakinolide (blue). *n* = 15/13/16/13, total SypHy puncta analysed were 341/382/430/311. **(C)** Quantitative analysis of endocytosis at 90 seconds after the end of stimulation (SypHy signal loss as % of exocytosis signal) in individual neurons (**left**), and means ± SEM (**right**). **(D)** Quantification of SypHy signal decay time constant (tau, monoexponential fit) in individual neurons (**left**), and means ± SEM (**right**). Statistical analysis was done using one-way ANOVA with Tukey *post hoc* test, ns: non-significant, **P< 0.01; *** P< 0.001.

### Release activity-dependent recruitment of N-cadherin to the peri-active zone of synapses

We further hypothesized that if N-cadherin is promoting vesicle membrane endocytosis, then it should be recruited to the endocytic/peri-active zone of synapses at high vesicle release activity to enhance the efficiency of vesicle endocytosis and thus ensure sufficient membrane retrieval. To study the localization of N-cadherin at synapses, we performed super-resolution fluorescence imaging using structured illumination microscopy (SIM) in cortical neurons in mass culture (12 DIV). To activate massive vesicle release, neurons were incubated in an extracellular soultion with high potassium ion concentration (50 mM) to induce strong depolarization of presynaptic release sites. We immunostained the cortical neurons for N-cadherin, vGlut1 (presynaptic marker) and PSD95 (postsynaptic marker) after 5 min K+ depolarization (or addition of control extracellular solution), and after an additional recovery period of 30 min in control extracellular solution, and performed SIM imaging of fluorescent puncta. First, we analysed K+ depolarization induced changes in N-cadherin and PSD95 puncta area. K+ depolarization led to a general increase in the area of N-cadherin puncta that tended to be reversible after the recovery period, whereas PSD95 puncta area was unchanged (Fig. 7A-C). This result confirmed a previous observation by Tanaka et al., 2000, that was described prior to the advent of super-resolution techniques, indicating activity-induced alterations in N-cadherin localization. To deeper investigate activity-induced changes in N-cadherin localization, we focused our analysis on synaptic sites, which were identified as co-localizations of vGlut1 and PSD95 (Fig. 7D-F). We found three categories of synapses in respect to N-cadherin localization: i) synapses with N-cadherin localized mainly at the peri-active zone (often assymetric), ii) synapses without detectable N-cadherin, and iii) synapses with N-cadherin localized exclusively at the synaptic cleft. Without K+ stimulation nearly 60% of synapses had cleft-associated, peri-active zone N-cadherin, about 40% of synapses had no detectable N-cadherin, and only few synapses (<5%) had exclusively synaptic cleft localized N-cadherin (Fig. 7G). Intriguingly, upon K+ depolarization we observed a significant, reversible increase in the fraction of synapses with cleft-associated, peri-active zone N-cadherin and a corresponding reversible decrease in the fraction of synapses without detectable N-cadherin. The fraction of synapses with exclusively synaptic cleft localized N-cadherin did not change significantly. These results suggest that N-cadherin gets recruited to the peri-active zone upon massive vesicle release, where it might strongly promote individual vesicle and bulk endocytosis to compensate for the release-associated membrane insertion (Fig. 8).

**Figure 7.**
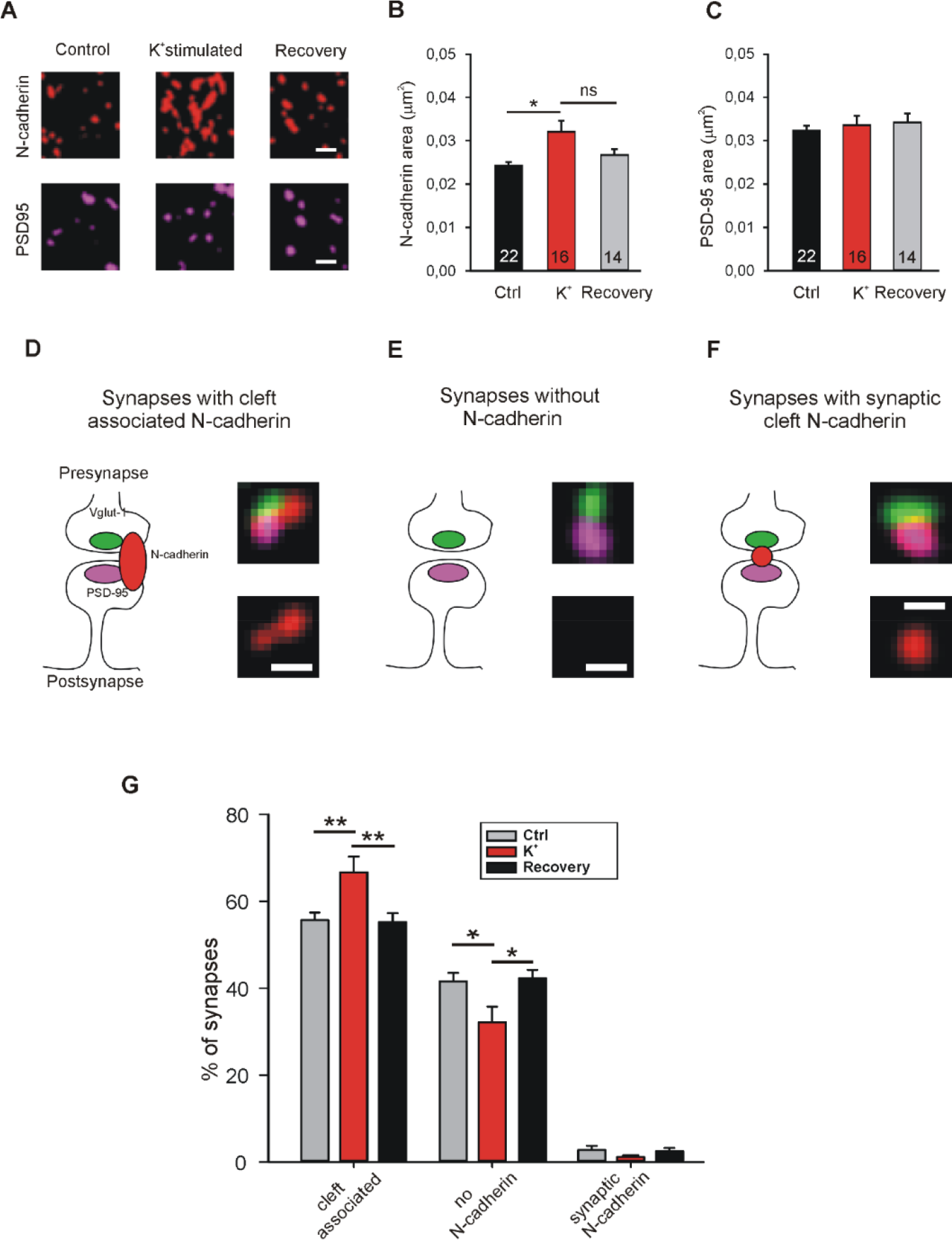
Recruitment of N-cadherin to glutamatergic synapses upon strong stimulation of vesicle release by high extracellular [K^+^]. **(A-C)** Structured illumination microscopy (SIM) revealed a significant increase in N-cadherin puncta area upon strong synaptic vesicle release induced by high extracellular [K^+^] (50 mM) in cultured cortical neurons (DIV 13-14). **(A)** Example images showing N-cadherin (upper panel) and PSD95 (lower panel) immunostaining in non-stimulated control cells (**left**), in 50 mM K^+^ stimulated cells (**middle**), and after 30 minutes recovery from K^+^ stimulation (**right**). Scale bar: 0.5 μm. **(B)** Quantification of mean N-cadherin puncta area (control: black; K^+^ stimulated: red; after recovery: grey). Means ± SEM, *n* = 22/16/14 dendrites (from four experiments). K^+^ stimulation resulted in a significant increase in N-cadherin puncta area**. (C)** Quantification of mean PSD95 puncta area. Statistics were done using Kruskal-Wallis one way ANOVA on Ranks with Dunn’s *post hoc* test, ns: non significant, * P< 0.05. **(D-F)** Schematic diagrams (**left**) and corresponding SIM overlay images (**right**) of glutamatergic synapses depicting three synapse categories as defined by the spatial expression of N-cadherin (red) in relation to VGLUT1 (presynaptic marker, green) and PSD95 (postsynaptic marker, magenta). **(D)** Synapses with cleft associated N-cadherin: N-cadherin is in close contact with synaptic cleft (at least at peri-active zone, **left**). Example SIM images showing VGLUT1, PSD95, N-cadherin signal overlay (**upper right**) and corresponding N-cadherin signal alone (**lower right**). **(E)** Synapses without N-cadherin expression (**left**). Example SIM images showing signal overlay (**upper right**) and absence of N-cadherin (**lower right**). **(F)** Synapses with synaptic cleft N-cadherin: N-cadherin expression exclusively inside the synaptic cleft (**left**). Example SIM images showing signal overlay (**upper right**) and N-cadherin signal alone (**lower right**). Only very few synapses had exclusively synaptic N-cadherin. Scale bar: 0.25 μm. **(G)** Quantification of the percentage of synapses in the three categories defined above (D-F) for non-stimulated controls (grey), in 50 mM K^+^ stimulated cells (red), and after 30 minutes recovery from K^+^ stimulation (black). Strong release of synaptic vesicles by K^+^ stimulation led to a significant increase in the number of synapses with cleft associated N-cadherin. *n* = 12 dendrites (from 4 experiments). Statistical analysis was done using one way ANOVA with Tukey *post hoc* test, * P< 0.05; ** P< 0.01.

**Figure 8.**
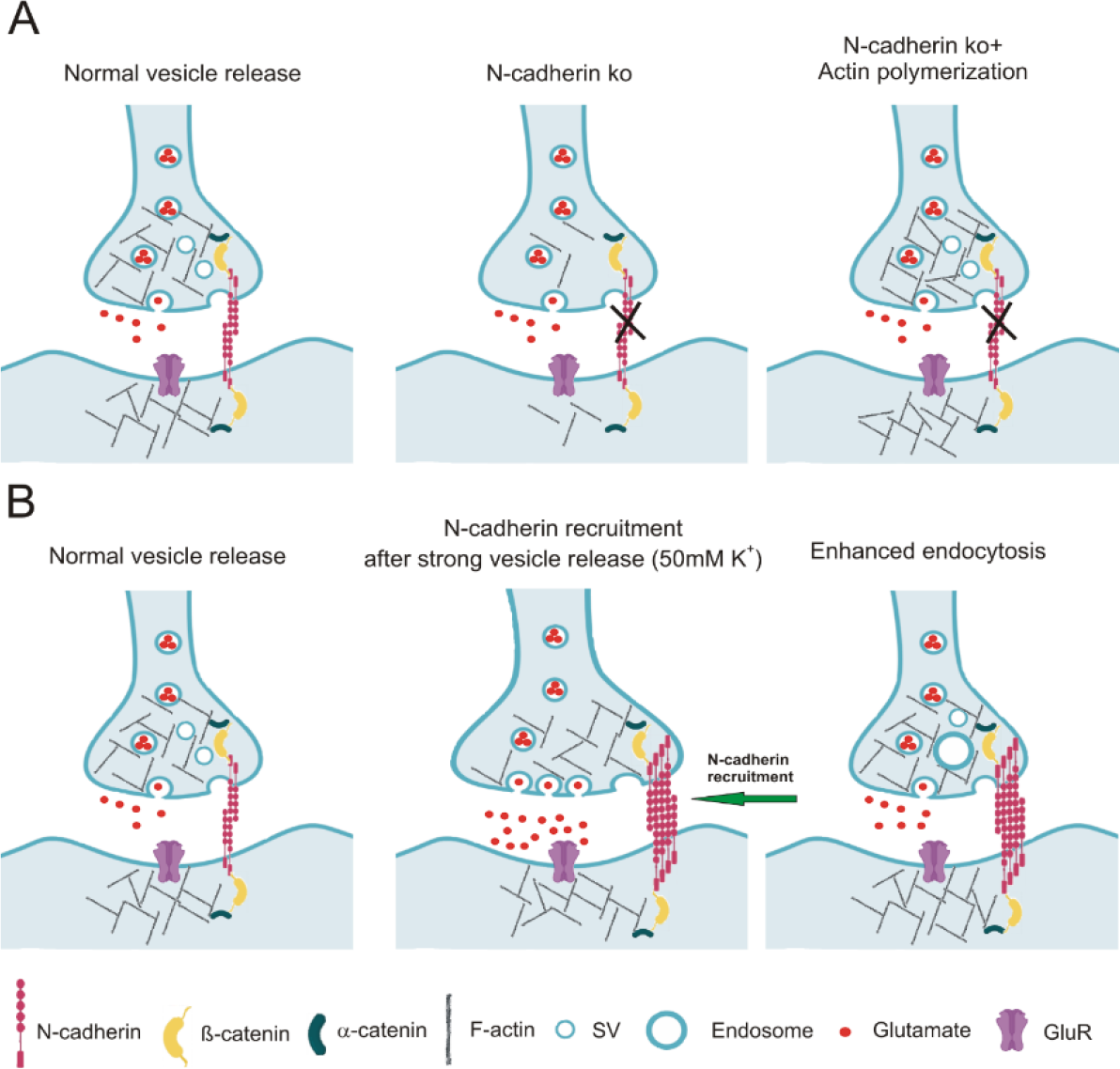
Model of the regulation of synaptic vesicle endocytosis by N-cadherin. (**A**) At normal vesicle release, loss of N-cadherin at the peri-active zone of glutamatergic synapses leads to disorganized actin filaments. This in turn results in reduced synaptic vesicle endocytosis. Impaired endocytosis is rescued by pharmacologically enhancing actin polymerization. (**B**) At massive vesicle release induced by K^+^ depolarization, additional N-cadherin is recruited to the peri-active zone of glutamatergic synapses. Enhanced N-cadherin clustering promotes actin filament organization, thus leading to more efficient endocytosis.

## Discussion

In this paper, our novel approach towards understanding the role of the synaptic adhesion molecule N-cadherin in synaptic vesicle endocytosis was to investigate the functional consequences of N-cadherin gene knockout at near physiological temperature. This turned out to be of crucial importance, because synaptic vesicle endocytosis is well known to be highly temperature sensitive (e.g. Granseth and Lagnado, 2008; Soykan et al., 2016; Chanaday and Kavalali, 2018). Using synaptophysin-pHluorin (SypHy) fluorescence imaging (Granseth et al., 2006; Royle et al., 2008; Kavalali and Jorgensen, 2014) in cultured cortical neurons, we were able to study both synaptic vesicle exo- and endocytosis in close temporal association to each other. At both, autaptic and axonal release sites, synaptic vesicle endocytosis was strongly enhanced regarding the amount and the kinetics of vesicle endocytosis upon increasing temperature from 25°C to 34°C. With moderate stimulation, at near physiological temperature a significant reduction and slow down of vesicle endocytosis was clearly evident at both autaptic and axonal release sites in N-cadherin knockout neurons. However, at room temperature no significant impairment of vesicle endocytosis was observed, possibly explaining why previous studies described mainly defects in vesicle exocytosis upon inhibiting N-cadherin function (Jüngling et al., 2006; Vitureira et al., 2011). This impairment of vesicle endocytosis in N-cadherin deficient neurons affected different modes of endocytosis similarly, thus resulting in altered single vesicle as well as bulk endocytosis. The defects in vesicle endocytosis observed in N-cadherin knockout neurons were largely confirmed in experiments, in which N-cadherin function was inhibited by overexpression of a dominant-negative C-terminal fragment that lacked the extracellular cadherin domains (NcadΔE). In addition to strong defects in vesicle endocytosis, we also found more difficult to detect defects in vesicle exocytosis in N-cadherin knockout neurons at both autaptic and axonal release sites. This required specific experimental paradigms that involved blockade of reacidification of vesicles with bafilomycin.

We further addressed for the first time the molecular mechanisms involved in the boosting of vesicle endocytosis by N-cadherin. We found that pharmacologically enhancing actin polymerization was able to rescue the defects in vesicle endocytosis in N-cadherin deficient neurons. This suggests that N-cadherin modulates endocytosis via its signaling to the actin filaments (Arrikath and Reichardt, 2008; Hirano and Takeichi, 2012) that are crucial for the endocytosis processes at synapses (Wu et al., 2016; Soykan et al., 2017). In a second step, we addressed whether N-cadherin might be involved in the compensatory enhancement of vesicle endocytosis that occurs upon strong vesicie release. Using super-resolution imaging, we observed that massive vesicle release leads to additional recruitment of N-cadherin to the peri-active zone, where it is in an optimal strategic position to enhance vesicle endocytosis via signaling to actin filaments (Fig. 8).

It has been well described by several recent studies and confirmed in this paper, that synaptic vesicle endocytosis is a highly temperature dependent process (e.g. Renden and Gersdorff, 2007; Watanabe et al., 2014; Delvendal et al., 2016). The temperature sensitivity of vesicle endocytosis is much more pronounced than the temperature dependence of vesicle exocytosis (Chanaday and Kavalali, 2018). This might be caused by the much more complex molecular mechanisms involved in vesicle endocytosis as compared to the comparatively simple molecular mechanism of vesicle exocytosis involving only a few crucial proteins. Therefore, gene knockout induced changes in vesicle exocytosis can well be studied at room temperature, which is used in the majority of papers on the function of specific proteins in exocytosis. In contrast, gene knockout studies on vesicle endocytosis at room temperature may be misleading, because even strong knockout effects might be obscured when vesicle endocytosis is analysed at very low efficiency and speed. It is therefore absolutely needed to analyse specific gene knockout effects on vesicle endocytosis at near physiological temperature. In this paper, we present to the best of our knowledge for the first time a detailed analysis of the effects of N-cadherin knockout on vesicle endocytosis at near physiological temperature in cortical synapses. We demonstrate that using near physiological temperature is crucial for clearly detecting significant effects of N-cadherin deficiency on synaptic vesicle endocytosis.

In this study, we investigated the effects of N-cadherin knockout on synaptic vesicle exo- and endocytosis at two rather different types of release sites, autapses and axonal release sites using SypHy imaging. Autapses in microisland cultures are well known to be very similar to *bona fide* synapses (Bekkers, 2020) with distinct presynaptic vesicle pools, i.e. a readily releasable (docked) pool, a reserve pool, and a resting pool (e.g. Rosenmund and Stevens, 1996; Mohrmann et al., 2003). Because of this large overall vesicle accumulation, it is difficult with standard stimulation and SypHy imaging to obtain a mean exocytosis signal of more than 40% of the NH_4_Cl signal (SypHy signal from all vesicles) at autapses. In contrast, axonal release sites do not contain large reserve and resting vesicle pools (Verderio et al., 1999; Matteoli et al., 2004). In line with this, we observed that the normalized SypHy exocytosis signal at axonal release sites was always larger, because the readily releasable pool represents a higher fraction of the total vesicle pool as compared to autapses. Autapses and axonal release sites also differ in respect to the consequences of the gene knockout for N-cadherin protein deficiency. At autapses N-cadherin is absent pre- and postsynaptically, while at axonal release sites N-cadherin is primarily absent only presynaptically. However, if a postsynaptic structure is present, presynaptic loss of N-cadherin might also induce a postsynaptic loss because of the transsynaptic homophilic interaction of N-cadherin. Despite these potential molecular differences between autapses and axonal release sites, we found that N-cadherin knockout resulted in similar defects in vesicle endocytosis as well as in vesicle exocytosis.

Synaptic vesicle endocytosis is well known to occur via several different modes with different kinetics (Clayton and Cousin, 2009a; Soykan et al., 2016; Soykan et al., 2017; Chananday and Kavalali, 2018; Gan and Watanabe, 2018). Depending on the exact experimental conditions slow clathrin-dependent individual vesicle endocytosis, fast kiss-and-run endocytosis, ultrafast endocytosis, and bulk endocytosis have been described. SypHy imaging appears to report well the more slow modes of endocytosis, i.e. individual vesicle endocytosis and bulk endocytosis. Here, we attempted to further distinguish between individual vesicle endocytosis and bulk endocytosis by studying the uptake of different fluorescent dyes (Clayton and Cousin, 2009b). These two modes were well separated by the specific experimental conditions and by the differential analysis used. Analysis of individual vesicle endocytosis confirmed an impairment of this mode of endocytosis in N-cadherin knockout neurons. In addition, analysis of bulk endocytosis revealed a similar impairment in N-cadherin deficient neurons suggesting that vesicle endocytosis in general might require the presence of N-cadherin to show optimal efficiency. In these experiments N-cadherin gene knockout was induced only in the postsynaptic neurons, because we did a sparse cre transfection of cortical neurons in mass culture. This indicates that a postsynaptic N-cadherin deficiency is sufficient to obtain a similar impairment of vesicle endocytosis as occuring with both pre- and postsynaptic knockout (autapses). This might be explained by the transsynaptic homophilic interaction of pre- and postsynaptic N-cadherin enabling a complete loss of function of the molecular N-cadherin adhesion system with only a postsynaptic knockout. In line with this idea, we showed in a previous paper (Jüngling et al., 2006) that postsynaptic N-cadherin knockout is sufficient to observe similar defects (on vesicle exocytosis) as obtained with both pre- and postsynaptic knockout.

In addition to inhibiting N-cadherin function by a gene knockout approach, we used overexpression of a dominant-negative C-terminal fragment of N-cadherin (NcadΔE) lacking the extracellular cadherin domains as an alternative approach to block N-cadherin function. The latter type of approach has been well established in the N-cadherin field by several studies (e.g. Togashi et al., 2002; Vitureira et al., 2011; Andreyeva et al., 2012). Using SypHy imaging at autapses in cultured cortical neurons, we observed very similar defects of NcadΔE expression as found with N-cadherin gene knockout strongly confirming an imporant role of N-cadherin in vesicle endocytosis. Interestingly, studying defects in endocytosis induced by postsynaptic NcadΔE expression by the uptake of fluorescent dyes revealed a clear impairment only for bulk endocytosis. Individual vesicle endocytosis directly from the plasma membrane appeared to be not affected by postsynaptic NcadΔE expression (see Fig. 4). This finding indicates that inhibition of N-cadherin function with NcadΔE overexpression is weaker than functional inhibition induced by N-cadherin gene knockout. This further suggests that bulk endocytosis might be more sensitive against partial loss of N-cadherin function possibly by its more complex nature involving large membrane invaginations as compared to individual vesicle endocytosis.

Regarding the molecular mechanisms of action of N-cadherin on vesicle endocytosis, the in general well known signaling of the N-cadherin adhesion complex from N-cadherin via beta-catenin and alpha-catenin to actin filaments (Arrikath and Reichardt, 2008; Hirano and Takeichi, 2012) is likely to be involved. It has been described recently, that vesicle endocytosis is strongly dependent on the state of actin filaments and is enhanced in the presence of actin polymerizing drugs (Soykan et al., 2017). In line with this mechanistic idea, we demonstrated here that the actin polymerizing drug jasplakinolide rescues the impairment of vesicle endocytosis that is induced by N-cadherin deficiency. Thus a modulatory influence of the N-cadherin adhesion complex on vesicle endocytosis via its classical signaling pathway to actin filaments appears conceivable.

Endocytosis at the peri-active zone has two basic functions: i) fast homeostatic membrane removal from the plasma membrane to compensate for the membrane addition that is occuring during vesicle exocytosis, and ii) retrieval of synaptic vesicles to maintain the reserve/resting pool, which is at hippocampal/cortical synapses rather large and therefore does not require fast refill. To enable fast homeostatic membrane removal, it appears to be needed that membrane endocytosis is enhanced upon massive vesicle release. Given the endocytosis enhancing action of N-cadherin demonstrated in this paper, a vesicle release-induced recruitment of N-cadherin to the peri-active zone might provide a molecular regulatory mechanism for fast homeostatic membrane removal by bulk endocytosis. To investigate a release activity-dependent recruitment of N-cadherin to the peri-active zone, we studied the synaptic localization of N-cadherin using super-resolution imaging (structured illumination microscopy). In line with previous reports (Uchida et al., 1996; Elste and Benson, 2006; Yam et al., 2013), we found that N-cadherin is localized most prominently at the peri-active zone of glutamatergic synapses, and also showed a somewhat weaker expression within the synaptic cleft. Similar to Yam et al., 2013, we unexpectedly observed a rather assymetric accumulation of N-cadherin mainly on one side of the synapse. Upon stimulating massive synaptic vesicle release with K+ depolarization, we observed a reversible, general increase in N-cadherin puncta as described previously (Tanaka et al., 2000; Yam et al., 2013). By further focussing the image analysis to *bona fide* glutamatergic synapses (identified by apposition of pre- and postsynaptic markers), we in addition observed a recruitment of N-cadherin to the peri-active zone of synapses upon K+ induced massive vesicle release. This release-induced recruitment of N-cadherin strongly supports a regulatory role of N-cadherin in homeostatic membrane removal (Fig. 8).

Previous studies focussing on the analysis of synaptic vesicle exocytosis in N-cadherin knockout ES cell-derived neurons (Jüngling et al., 2006), or using expression of a dominant-negative N-cadherin fragment in hippocampal neurons (Vitureira et al., 2011) described slight deficits in vesicle release. These defects were only detectable upon stimulating vesicle release with repetitive action potentials indicating that the refill of the readily releasable vesicle pool was impaired (Jüngling et al., 2006; Vitureira et al., 2011). Here, we confirmed such defects in vesicle exocytosis in N-cadherin knockout cortical neurons using repetitive stimulation at 20 Hz with SypHy imaging at both autaptic and axonal release sites. Regarding the molecular mechanisms involved, two working hypotheses appear well conceivable. Loss of N-cadherin function might via dysregulation of actin filament organization lead to an impaired movement of vesicles from the reserve pool to the docked vesicle pool. Alternatively, loss of N-cadherin function might again via dysregulation of actin filaments result in an impairment of release site clearance thus inhibiting further vesicle release.

## Conclusions

In addition to its well established postsynaptic role in spine plasticity, the transsynaptic N-cadherin adhesion complex has an important presynaptic regulatory function in synaptic vesicle endocytosis as indicated by our N-cadherin knockout study in cortical neurons (Fig. 8A). The endocytosis enhancing action of N-cadherin appeared to be mediated by promoting presynaptic actin filament organization. Because N-cadherin was recruited to the peri-active zone of glutamatergic synapses upon massive vesicle release (Fig. 8B), its endocytosis boosting effect might lead to enhanced membrane endocytosis thus compensating for the vesicle release-induced enlargement of the presynaptic plasma membrane.

## Materials and Methods

Methods were carried out in accordance with all relevant guidelines.

### Cell culture

Primary neuronal mass cultures were obtained from cortices of C57/BL6 wildtype mice, and from Ncad^flox/flox^ mice (Kostetskii et al., 2005) as described previously (van Stegen et al., 2017; Dagar et al., 2019). In brief, cortices from E18 fetuses were dissected into small pieces and were treated with trypsin for 5 min. These pieces were mechanically triturated in Basal Medium Eagle (BME) media (Gibco) after removal of trypsin. Dissociated neurons were pelleted down by centrifugation, resuspended, and 20,000-30,000 neurons were seeded on poly-L-ornithine coated glass cover slips. Fresh Neurobasal (NB) medium (Gibco) with B27 supplement (2%, 50x; Gibco) containing penicillin-streptomycin (Gibco) and Glutamax-1 (Gibco) was added to the cells after 1-2 h. These mass cultures were further cultivated and maintained in a humidified incubator with 5% CO_2_ at 37°C for 12-15 days *in vitro* (DIV).

Autaptic glial microisland cultures were done as described previously (Mohrmann et al., 2003; van Stegen et al., 2017). In brief, mouse astrocytes were obtained by dissociating cortices of P0-P2 pups from C57/BL6 wildtype mice, and culturing the cells in BME medium for 10-12 days to form a confluent monolayer. These astrocytes were then re-dissociated, seeded at low density on glass coverslips, and further cultured for 5-7 days in BME medium to form glial microislands. Then, 20.000–30.000 dissociated cortical neurons (see above) were seeded per coverslip. After 5-6 hours, BME medium was replaced by NB medium containing B27 supplement, penicillin-streptomycin (Gibco) and Glutamax-1 (Gibco). The culture medium was exchanged every 3 days during further cultivation (12-16 DIV).

### Transfection and plasmids

Cultures were transfected by using magnetic nanobeads (NeuroMag; OZ Biosciences) as described previously (van Stegen et al., 2017; Dagar et al., 2019). Briefly, plasmid DNA and Neuromag were mixed in NB medium without any supplement, and were incubated for 20 min at room temperature to form complexes. These complexes were then directly added to the neurons. To enhance transfection efficiency, an oscillating magnetic field was applied for 30 min by placing the cultured neurons (6-well plate) on a magnetic plate (Magnetofection™, magnefect LT; nanoTherics). After transfection, 500 μl of fresh culture medium was added. The following plasmids were used: EBFP2-N1 (a gift from Dr. Michael Davidson, Florida, USA; addgene #54595), pDsRed2-N1 (Clontech), SypHy-A4 (gift from Dr. L. Lagnado, Cambridge, UK, addgene #24478), pBS598EF1alpha-EGFPcre (addgene), and pcDNA3.1-FLAG-NCadCTF1 (ΔE N-cadherin expression plasmid; Andreyeva et al., 2012).

### Wide-field fluorescence microscopy and electrical stimulation

Wide-field fluorescence imaging was performed using an inverted motorized Axiovert 200 M microscope (Zeiss) with a 12 bit CoolSnap ES2 CCD camera (Photometrics) and 40x/1.3 oil objective using MetaVue or Visiview software (Molecular Devices/Visitron Systems) unless mentioned. The following filter sets (Zeiss) were used: 1) for SypHy and FM1-43: excitation 485/20 nm, beam splitter 510 nm, emission 515/565 nm; 2) for DsRed2 and TM-dextran: excitation 545/25 nm, beam splitter 570 nm, emission 605/70. To induce synaptic vesicle exo- and endocytosis by electrical stimulation (for SypHy imaging, and for FM1-43 / TMR-dextran imaging), cortical neurons were mounted in a stimulation chamber (Chamlide, Live Cell Instrument, Republic of Korea) on the stage of an Axiovert 200 M microscope. Extracellular field stimulation with 1 ms biphasic pulses of 100 mA (A385/A382 World Precision Instruments) at 20 Hz was used.

### Synaptophysin-pHluorin (SypHy) imaging and data analysis

To study synaptic vesicle exocytosis and endocytosis, Synaptophysin-pHluorin (SypHy) imaging was performed according to standard protocols (Granseth et al., 2006; Royle et al., 2008). In brief, autaptic microisland cultures were co-transfected with SypHy-A4 and DsRed2 (and CreEGFP or NcadΔE), and SypHy experiments were performed 3 to 5 days post-transfection. The transfected cultures were mounted in a stimulation chamber (Live Cell Instrument) onto the stage of an Axiovert 200 M microscope (Zeiss), and were maintained in extracellular solution (in mM: 136 NaCl, 2.5 KCl, 2 CaCl_2_, 1.3 MgCl_2_, 10 HEPES, 10 glucose, pH = 7.3). DL-AP5 (50 μM) and DNQX (10 μM) were added to the extracellular medium to prevent recurrent network activity. Synaptic vesicle cycling was induced by electrical stimulation. Fluorescence images were taken at time intervals of 2 seconds, and in total a 4 min image sequence was done to record the stimulation-induced changes in SypHy fluorescence. At the end of each experiment an extracellular solution containing a high concentration of NH_4_Cl (in mM: 50 NH_4_Cl, 86 NaCl, 2.5 KCl, 2 CaCl_2_, 1.3 MgCl_2_, 10 HEPES, 10 glucose, pH = 7.3) was added to the cells to alkalize the total vesicle pool. This resulted in a maximal SypHy fluorescence signal that was used for normalization required for quantitative data analysis.

SypHy fluorescence intensities over time were analysed offline using Metamorph software. For this, DsRed2 images were overlayed on maximum SypHy fluorescence images obtained by adding high NH_4_Cl containing extracellular solution at the end of each experiment. Regions of interest (ROIs) were defined around SypHy puncta located on dendrites of the transfected neuron for autaptic contacts, and around SypHy puncta not contacting any dendrite of the transfected neuron for axonal release sites. These ROIs were transferred to the fluorescence image sequence (including response to electrical stimulation) to obtain average SypHy fluorescence intensities over time for each punctum, and these intensity values were transferred to Microsoft excel software for further analysis. The average fluorescence intensity within a ROI was first background subtracted at each time point, and then the baseline fluorescence intensity (value of time point just prior to begin of electrical stimulation) was subtracted to compensate for differences in baseline fluorescence. (For figure 5, the average of ten successive baseline values before the begining of electrical stimulation was used to subtract baseline intensity.) Finally, these intensity values were normalized to the maximal fluorescence signal (background subtracted) obtained at the same punctum after adding NH_4_Cl containing extracellular solution at the end of each experiment. SypHy puncta not showing a fluorescence increase upon stimulation (non-releasing sites) were excluded from analysis.

### FM1-43 and TMR-dextran co-staining and data analysis

To differentially label synaptic vesicle endocytosis occurring by different mechanisms, cultured neurons were co-stained by using the styryl dye FM1-43 and Tetramethylrhodamine(TMR)–dextran at room (25°C) and at near physiological temperature (34°C) (Clayton and Cousin, 2009b). Cultured neurons were mounted in a stimulation chamber (Live Cell Instrument) and synaptic vesicle turnover was induced by electrical stimulation (at 20 Hz). TMR–dextran (40 kDa, 50 μM) and FM1-43 (10 μM) were present during the stimulation, and were removed after the end of stimulation by immediate washing for 3-5 min in Ca^2+^-free extracellular solution containing 1 mM ADVASEP-7. To ensure the specificity of TMR-dextran uptake, quantitative analysis was done only for TMR-dextran puncta colocalizing with FM1-43 puncta. For quantitative analysis of FM1-43 and TMR-dextran co-stainings, fluorescence images were thresholded offline using MetaMorph software (Molecular Devices/Visitron Systems), and a low pass filter was applied to remove single pixel noise. FM1-43 and TMR-dextran fluorescence images were then overlayed on the images of EBFP2 expressing neurons to identify FM1-43 puncta on dendrites of transfected neurons. Regions of interest (ROI) were created around FM1-43 puncta to quantify the puncta density / 10 μm of dendrite length (total endocytosis signal). To identify the bulk endocytosis component, the density of FM1-43 puncta co-localizing with TMR-dextran on the dendrite of transfected neurons was determined (bulk endocytosis signal).

### Stimulation by high extracellular [K^+^] and immunocytochemistry

To study the structural changes associated with strong synaptic vesicle release, cultured cortical neurons (12-13 DIV) were stimulated for 5 minutes by using an extracellular solution containing high [K^+^] (in mM: 88.5 NaCl, 50 KCl, 2 CaCl_2_, 1.3 MgCl_2_, 10 HEPES, 10 glucose, pH = 7.3). After stimulation, cultures were immediately fixed with 4% paraformaldehyde (PFA). For control, cultured neurons were incubated in a standard extracellular solution (in mM: 136 NaCl, 2.5 KCl, 2 CaCl_2_, 1.3 MgCl_2_, 10 HEPES, 10 glucose, pH = 7.3) for 5 minutes before fixation. In addition, neuronal cultures were allowed to recover for 30 minutes in a standard extracellular solution (2.5 mM [K^+^]) after stimulation (5 minutes in 50 mM [K^+^]), and then fixed with 4% PFA (20 min). After stimulation and fixation, cells were subjected to a triple immunocytochemical staining according to a standard protocol. In brief, cells were permeabilized with 0.3% Triton X-100 in blocking buffer (10% fetal bovine serum (FBS), 5% sucrose, 2% bovine serum albumin (BSA), in PBS, pH 7.4) for 30 min at room temperature. Cells were then incubated for 1 h with primary antibodies, followed by washing 3×10 min with phosphate buffered saline (PBS). After this, cultures were incubated with Alexa Fluor (AF) conjugated secondary antibodies (Invitrogen) for 1 h at room temperature. Then, cultures were again rinsed 3×10 min with PBS, and coverslips were mounted to reduce photobleaching. The following primary antibodies were used: anti-N-Cadherin (rabbit polyclonal, 1:500, Abcam; Cat. No. 18203), anti-VGLUT1 (guinea pig polyclonal, 1:500, Synaptic Systems Cat.No.135304), and anti-PSD95 (mouse monoclonal, 1:500, Thermo Fisher Scientific MA1-046). The following secondary antibodies were used: AF 555 (goat anti-rabbit, 1:500, Cat. No. A21429), AF488 (goat anti-guinea pig, 1:500, Cat. No. A11073), and AF647 (goat anti-mouse, 1:500, Cat. No. A21241).

### Structured illumination microscopy (SIM) imaging and data analysis

Super-resolution structured illumination microscopy (SIM) imaging was performed using an ELYRA PS microscope (Zeiss) at the Center for Advanced Imaging (CAI), HHU Düsseldorf. Three different diode lasers with 488 nm, 561 nm, and 642 nm wavelengths were used to excite different fluorophores in triple immunostained cultured cortical neurons (see above). Images were captured with an alpha Plan-Apochromat 63x/1.4 oil DIC M27 objective (Zeiss) as z-stacks using an EM-CCD camera with 1002 x 1002 pixels for maximum resolution. To obtain super-resolution, out of focus light was computationally removed (optical sectioning) and super-resolution images were reconstructed offline. Maximum intensity projections were used for further analysis. First, images (individual channels) were thresholded based on the intensity profile using ImageJ Fiji, and single pixel noise was removed using a median filter. The thresholded images were given pseudo colours, and an overlay was created using VGLUT1 and PSD95 images. The overlay was then searched for co-localizing VGLUT1+PSD95 puncta (glutamatergic synapses), and regions of interest were created around them. The corresponding N-cadherin image was then superimposed on the overlay, and N-cadherin localization was determined in relation to synapses (VGLUT1+PSD95 puncta).

### Statistics

All data are given as individual values and as means ± SEM. Statistical significance was determined by Student’s t-test, if applicable (whenever normality test failed, Mann-Whitney’s test on ranks was performed), and by one-way ANOVA in combination with Tukey’s posthoc test. SigmaPlot 11 software was used.

## Acknowledgements

We want to thank Dr. B. van Stegen, Heinrich-Heine University Düsseldorf for introduction into SypHy imaging, and Dr. L. Lagnado, Dr. M. Davidson, and Dr. A. Andreyeva for providing expression vectors. We further want to thank Dr. H. Aberle and Rebekka Ochs, Heinrich-Heine University Düsseldorf, for introduction into the use of the Zeiss ELYRA PS SIM microscope, that was provided by the Center for Advanced Imaging (CAI) of the Heinrich-Heine University Düsseldorf. We further want to thank Dr. B. van Stegen for performing initial experiments with SIM imaging.

## Author contributions

SD and KG designed research, SD performed experiments and analysed data, SD and KG wrote the paper.

## References

Abe, K., Chisaka, O., Van Roy, F., and Takeichi, M. (2004). Stability of dendritic spines and synaptic contacts is controled by alpha N-catenin. Nat Neurosci 7, 357–363.

Andreyeva, A., Nieweg, K., Horstmann, K., Klapper, S., Muller-Schiffmann, A., Korth, C., and Gottmann, K. (2012). C-terminal fragment of N-cadherin accelerates synapse destabilization by amyloid-beta. Brain 135, 2140–2154.

Arikkath, J., and Reichardt, L.F. (2008). Cadherins and catenins at synapses: roles in synaptogenesis and synaptic plasticity. Trends Neurosci 31, 487–494.

Atluri, P.P., and Ryan, T.A. (2006). The kinetics of synaptic vesicle reacidification at hippocampal nerve terminals. J Neurosci 26, 2313–2320.

Bamji, S.X., Shimazu, K., Kimes, N., Huelsken, J., Birchmeier, W., Lu, B., and Reichardt, L.F. (2003). Role of beta-catenin in synaptic vesicle localization and presynaptic assembly. Neuron 40, 719–731.

Bekkers, J.M. (2020). Autaptic cultures: methods and applications. Front Synaptic Neurosci 12, 18.

Benson, D.L., and Huntley, G.W. (2012). Synapse adhesion: a dynamic equilibrium conferring stability and flexibility. Curr Opin Neurobiol 22, 397–404.

Bian, W.J., Miao, W.Y., He, S.J., Qiu, Z., and Yu, X. (2015). Coordinated spine pruning and maturation mediated by inter-spine competition for cadherin/catenin complexes. Cell 162, 808–822.

Bozdagi, O., Wang, X.B., Nikitczuk, J.S., Anderson, T.R., Bloss, E.B., Radice, G.L., Zhou, Q., Benson, D.L., and Huntley, G.W. (2010). Persistence of coordinated long-term potentiation and dendritic spine enlargement at mature hippocampal CA1 synapses requires N-cadherin. J Neurosci 30, 9984–9989.

Brasch, J., Harrison, O.J., Honig, B., and Shapiro, L. (2012). Thinking outside the cell: how cadherins drive adhesion. Trends Cell Biol 22, 299–310.

Brockhaus, J., Schreitmuller, M., Repetto, D., Klatt, O., Reissner, C., Elmslie, K., Heine, M., and Missler, M. (2018). alpha-Neurexins together with alpha2delta-1 auxiliary subunits regulate Ca2+ influx through Cav2.1 channels. J Neurosci 38, 8277–8294.

Chanaday, N.L., and Kavalali, E.T. (2018). Time course and temperature dependence of synaptic vesicle endocytosis. FEBS Lett 592, 3606–3614.

Chen, C.Y., Chen, Y.T., Wang, J.Y., Huang, Y.S., and Tai, C.Y. (2017). Postsynaptic Y654 dephosphorylation of beta-catenin modulates presynaptic vesicle turnover through increased N-cadherin mediated transsynaptic signaling. Dev Neurobiol 77, 61–74.

Clayton, E.L., and Cousin, M.A. (2009a). The molecular physiology of activity-dependent bulk endocytosis of synaptic vesicles. J Neurochem 111, 901–914.

Clayton, E.L., and Cousin, M.A. (2009b). Quantitative monitoring of activity-dependent bulk endocytosis of synaptic vesicle membrane by fluorescent dextran imaging. J Neurosci Methods 185, 76–81.

Dagar, S., and Gottmann, K. (2019). Differential properties of the synaptogenic actvities of the neurexin ligands neuroligin1 and LRRTM2. Front Mol Neurosci 12, 269.

Delvendahl, I., Vyleta, N.P., von Gersdorff, H., and Hallermann, S. (2016). Fast, temperature-sensitive and clathrin-independent endocytosis at central synapses. Neuron 90, 492–498.

Elste, A.M., and Benson, D.L. (2006). Structural basis for developmentally regulated changes in cadherin function at synapses. J Comp Neurol 495, 324–335.

Friedman, L.G., Benson, D.L., and Huntley, G.W. (2015). Cadherin-based transsynaptic networks in establishing and modifying neural connectivity. Curr Top Dev Biol 112, 415–465.

Gaffield, M.A., and Betz, W.J. (2006). Imaging synaptic vesicle exocytosis and endocytosis with FM dyes. Nat Protoc 1, 2916–2921.

Gan, Q., and Watanabe, S. (2018). Synaptic Vesicle Endocytosis in Different Model Systems. Front Cell Neurosci 12, 171.

Granseth, B., Odermatt, B., Royle, S.J., and Lagnado, L. (2006). Clathrin-mediated endocytosis is the dominant mechanism of vesicle retrieval at hippocampal synapses. Neuron 51, 773–786.

Granseth, B., and Lagnado, L. (2008). The role of endocytosis in regulating the strength of hippocampal synapses. J Physiol 586, 5969–5982.

Han, K.A., Ko, J.S., Pramanik, G., Kim, J.Y., Tabuchi, K., Um, J.W., and Ko, J. (2018). PTPsigma drives excitatory presynaptic assembly via various extracellular and intracellular mechanisms. J Neurosci 38, 6700–6721.

Hirano, S., and Takeichi, M. (2012). Cadherins in brain morphogenesis and wiring. Physiol Rev 92, 597–634.

Hua, Y., Sinha, R., Thiel, C.S., Schmidt, R., Huve, J., Martens, H., Hell, S.W., Egner, A., and Klingauf, J. (2011). A readily retrievable pool of synaptic vesicles. Nat Neurosci 14, 833–839.

Jüngling, K., Eulenburg, V., Moore, R., Kemler, R., Lessmann, V., and Gottmann, K. (2006). N-cadherin transsynaptically regulates short-term plasticity at glutamatergic synapses in embryonic stem cell-derived neurons. J Neurosci 26, 6968–6978.

Kadowaki, M., Nakamura, S., Machon, O., Krauss, S., Radice, G.L., and Takeichi, M. (2007). N-cadherin mediates cortical organization in the mouse brain. Dev Biol 304, 22–33.

Kavalali, E.T., and Jorgensen, E.M. (2014). Visualizing presynaptic function. Nat Neurosci 17, 10–16.

Kostetskii, I., Li, J., Xiong, Y., Zhou, R., Ferrari, V.A., Patel, V.V., Molkentin, J.D., and Radice, G.L. (2005). Induced deletion of the N-cadherin gene in the heart leads to dissolution of the intercalated disc structure. Circ Res 96, 346–354.

Li, M.Y., Miao, W.Y., Wu, Q.Z., He, S.J., Yan, G., Yang, Y., Liu, J.J., Taketo, M.M., and Yu, X. (2017). A critical role of presynaptic cadherin/catenin/p140cap complexes in stabilizing spines and functional synapses in the neocortex. Neuron 94, 1155–1172.

Matteoli, M., Coco, S., Schenk, U., and Verderio, C. (2004). Vesicle turnover in developing neurons: how to build a presynaptic terminal. Trends Cell Biol 14, 133–140.

Mendez, P., De Roo, M., Poglia, L., Klauser, P., and Muller, D. (2010). N-cadherin mediates plasticity-induced long-term spine stabilization. J Cell Biol 189, 589–600.

Mohrmann, R., Lessmann, V., and Gottmann, K. (2003). Developmental maturation of synaptic vesicle cycling as a distinctive feature of central glutamatergic synapses. Neuroscience 117, 7–18.

Murase, S., Mosser, E., and Schuman, E.M. (2002). Depolarization drives beta-Catenin into neuronal spines promoting changes in synaptic structure and function. Neuron 35, 91–105.

Mysore, S.P., Tai, C.Y., and Schuman, E.M. (2008). N-cadherin, spine dynamics, and synaptic function. Front Neurosci 2, 168–175.

Okuda, T., Yu, L.M., Cingolani, L.A., Kemler, R., and Goda, Y. (2007). beta-Catenin regulates excitatory postsynaptic strength at hippocampal synapses. Proc Natl Acad Sci USA 104, 13479–13484.

Radice, G.L., Rayburn, H., Matsunami, H., Knudsen, K.A., Takeichi, M., and Hynes, R.O. (1997). Developmental defects in mouse embryos lacking N-cadherin. Dev Biol 181, 64–78.

Renden, R., and von Gersdorff, H. (2007). Synaptic vesicle endocytosis at a CNS nerve terminal: faster kinetics at physiological temperatures and increased endocytotic capacity during maturation. J Neurophysiol 98, 3349–3359.

Rosenmund, C., and Stevens, C.F. (1996). Definition of the readily releasable pool of vesicles at hippocampal synapses. Neuron 16, 1197–1207.

Royle, S.J., Granseth, B., Odermatt, B., Derevier, A., and Lagnado, L. (2008). Imaging phluorin-based probes at hippocampal synapses. Methods Mol Biol 457, 293–303.

Sando, R., Jiang, X., and Sudhof, T.C. (2019). Latrophilin GPCRs direct synapse specificity by coincident binding of FLRTs and teneurins. Science 363, eaav7969.

Scheiffele, P., Fan, J., Choih, J., Fetter, R., and Serafini, T. (2000). Neuroligin expressed in nonneuronal cells triggers presynaptic development in contacting axons. Cell 101, 657–669.

Schreiner, D., Savas, J.N., Herzog, E., Brose, N., and de Wit, J. (2017). Synapse biology in the ‘circuit-age’-paths toward molecular connectomics. Curr Opin Neurobiol 42, 102–110.

Soykan, T., Kaempf, N., Sakaba, T., Vollweiter, D., Goerdeler, F., Puchkov, D., Kononenko, N.L., and Haucke, V. (2017). Synaptic vesicle endocytosis occurs on multiple timescales and is mediated by formin-dependent actin assembly. Neuron 93, 854–866.

Soykan, T., Maritzen, T., and Haucke, V. (2016). Modes and mechanisms of synaptic vesicle recycling. Curr Opin Neurobiol 39, 17–23.

Stan, A., Pielarski, K.N., Brigadski, T., Wittenmayer, N., Fedorchenko, O., Gohla, A., Lessmann, V., Dresbach, T., and Gottmann, K. (2010). Essential cooperation of N-cadherin and neuroligin-1 in the transsynaptic control of vesicle accumulation. Proc Natl Acad Sci USA 107, 11116–11121.

Südhof, T.C. (2017). Synaptic neurexin complexes: a molecular code for the logic of neural circuits. Cell 171, 745–769.

Südhof, T.C. (2018). Towards an understanding of synapse formation. Neuron 100, 276–293.

Takeichi, M. (2007). The cadherin superfamily in neuronal connections and interactions. Nat Rev Neurosci 8, 11–20.

Tanaka, H., Shan, W., Phillips, G.R., Arndt, K., Bozdagi, O., Shapiro, L., Huntley, G.W., Benson, D.L., and Colman, D.R. (2000). Molecular modification of N-cadherin in response to synaptic activity. Neuron 25, 93–107.

Togashi, H., Abe, K., Mizoguchi, A., Takaoka, K., Chisaka, O., and Takeichi, M. (2002). Cadherin regulates dendritic spine morphogenesis. Neuron 35, 77–89.

Uchida, N., Honjo, Y., Johnson, K.R., Wheelock, M.J., and Takeichi, M. (1996). The catenin/cadherin adhesion system is localized in synaptic junctions bordering transmitter release zones. J Cell Biol 135, 767–779.

van Stegen, B., Dagar, S., and Gottmann, K. (2017). Release activity-dependent control of vesicle endocytosis by the synaptic adhesion molecule N-cadherin. Sci Rep 7, 40865.

Verderio, C., Coco, S., Bacci, A., Rossetto, O., De Camilli, P., Montecucco, C., and Matteoli, M. (1999). Tetanus toxin blocks the exocytosis of synaptic vesicles clustered at synapses but not of synaptic vesicles in isolated axons. J Neurosci 19, 6723–6732.

Vitureira, N., Letellier, M., White, I.J., and Goda, Y. (2011). Differential control of presynaptic efficacy by postsynaptic N-cadherin and beta-catenin. Nat Neurosci 15, 81–89.

Watanabe, S., Trimbuch, T., Camacho-Perez, M., Rost, B.R., Brokowski, B., Sohl-Kielczynski, B., Felies, A., Davis, M.W., Rosenmund, C., and Jorgensen, E.M. (2014). Clathrin regenerates synaptic vesicles from endosomes. Nature 515, 228–233.

Wu, X.S., Lee, S.H., Sheng, J., Zhang, Z., Zhao, W.D., Wang, D., Jin, Y., Charnay, P., Ervasti, J.M., and Wu, L.G. (2016). Actin is crucial for all kinetically distinguishable forms of endocytosis at synapses. Neuron 92, 1020–1035.

Yam, P.T., Pincus, Z., Gupta, G.D., Bashkurov, M., Charron, F., Pelletier, L., and Colman, D.R. (2013). N-cadherin relocalizes from the periphery to the center of the synapse after transient synaptic stimulation in hippocampal neurons. PLoS One 8, e79679.

