## Supplementary material for "Transsynaptic N-cadherin adhesion complexes control presynaptic vesicle and bulk endocytosis at physiological temperature": Suppl. Figure 1

**Supplemental Material**

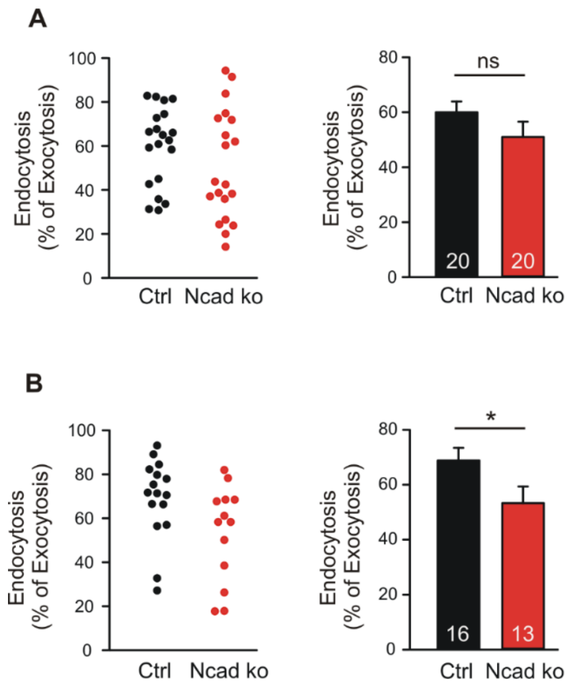


**Supplementary Figure 1. Reduced synaptic vesicle endocytosis was observed in N-cadherin knockout neurons using 100 stimuli at 20 Hz in SypHy imaging experiments** Quantitative analysis of synaptic vesicle endocytosis from the SypHy experiments (100 stimuli at 20 Hz at 34°C) shown in Fig. 5 A-D. (**A**) Quantitative analysis of endocytosis at 50 seconds after the end of stimulation (SypHy signal loss as % of exocytosis signal; values at 48, 50 and 52 sec averaged; 100 stimuli at 20 Hz) at autapses in individual neurons (**left**), and means ± SEM for control (black) and N-cadherin knockout (red) neurons (**right**). *n* = 20/20 cells. Student's *t*-test was used for statistical analysis, ns: non-significant. (**B**) Quantitative analysis of endocytosis at 50 seconds after the end of stimulation (SypHy signal loss as % of exocytosis signal; values at 48, 50 and 52 sec averaged; 100 stimuli at 20 Hz) at axonal release sites in individual neurons (**left**), and means ± SEM for control (black) and N-cadherin knockout (red) neurons (**right**). *n* = 16/13 cells. Mann-Whitney's test on ranks was performed for statistical analysis,* P<0.05.
